# The geometry of cortical sound processing in slow wave sleep

**DOI:** 10.1101/2025.06.13.658070

**Authors:** Allan Muller, Anton Filipchuk, Sophie Bagur, Brice Bathellier

## Abstract

During wake, sound-evoked and spontaneous neural activity of the auditory cortex evolve in distinct subspaces whereas anesthesia disrupts sound responses and merges these spaces. To evaluate if similar modifications of the sound representation geometry explain sensory disconnection during sleep, we followed large neural populations of the mouse auditory cortex across slow wave sleep and wakefulness. We observed that sleep dampens sound responses but preserves the geometry of sound representations which remain separate from spontaneous activity. Moreover, response dampening was strongly coordinated across neurons and varied throughout sleep spanning from fully preserved response patterns to population response failures on a fraction of sound presentations. These failures are more common during high spindle-band activity and more rarely observed in wakefulness. Therefore, in sleep, the auditory system preserves sound feature selectivity up to the cortex for detailed acoustic surveillance, but concurrently implements an intermittent gating mechanism leading to local sensory disconnections.

## Introduction

Sleep is characterized by reduced motor activity, decreased behavioral responsiveness and an absence of memorization of external stimuli which are often interpreted as a sensory disconnection of the brain from the environment^1–3^. Several neural mechanisms have been proposed to explain this disconnection, including a gating of sensory information by the thalamus based on its reduced activity, burst mode dynamics and the occurrence of spindles^4^. However, cortical sensory responses are still observed during sleep, in particular for its auditory part ^5,6^, even during spindles^7^ which invalidates this hypothesis. Moreover, the brain of sleeping subjects still specifically responds to complex stimuli, including the subject’s name^8^, extreme threats^9^, or learnt behavioral associations^10^. This remnant processing may preserve some environmental monitoring. At the same time, core sleep processes such as memory consolidation or network homeostasis appear to require isolation from external input in order to faithfully reflect internal constraints^11,12^. This raises the question of how the brain combines surveillance and offline processes in sleep.

Inspired by the recent observation that in spite of continued individual neuron responses, population-level sensory representations are deeply modified under anesthesia^13^, we reasoned that a specific re-organization of sensory representation geometry in sleep could subserve these two antagonist functions. We therefore followed the activity of up to a thousand neurons across wakefulness and sleep in the mouse auditory cortex during spontaneous activity and sound processing. We observed that sleep significantly reduces the magnitude of population responses to sounds and the amount of available sound information. However, contrary to anesthesia, this reduced responsiveness preserves the structure of sound representation and the separation of spontaneous and evoked activity subspaces. This indicates that sensory processing is intact, but dampened. Focusing on single trial population responses, we observed that this dampening is due to an adjustment of response gain, coordinated across local cortical populations, which intermittently leads to a full disconnection of the network from the incoming sound input. This intermittent switching between normal sensory processing and disconnection may allow combining frequent auditory monitoring and offline functions.

## Results

### Robust sleep during head-fixed two-photon calcium imaging

To follow spontaneous and sound-evoked activity across different sleep states and wakefulness, we trained mice to remain head-fixed on a running-wheel under a fully silent acousto-optic two-photon microscope (**Fig. 1a,b**). During the imaging session, we alternated 2 min periods of silence and 2 min periods of sound presentations (60 sounds, approximately 500 ms duration, inter-sound interval 2s) to sample spontaneous and sound-evoked activity. We imaged layer 2/3 neurons expressing GCAMP6s under the synapsin promoter using simultaneous acquisition of 4 planes at a global sampling rate of 23 Hz (**Fig. 1c**), yielding on average 649 ± 307 neurons per session. Calcium signals were extracted after motion correction^14^ and the calcium event detection algorithm MLSpike was used to denoise calcium signals into an event time series that approximates the actual spike trains of neurons^13,15^ (**Fig. 1d**). The sounds included pure tones, frequency chirps, tones with linear and sinusoidal amplitude modulations, broadband noises, and complex natural sounds (**Fig 1e**). Sleep scoring was performed in an automated manner based on local field potentials recorded in the dorsal hippocampus, the olfactory bulb and auditory cortex, contralateral to the imaging window, and on an electromyogram (EMG) electrode placed on the neck muscle, using previously established methodology^16^ (**Fig. 1a,f-i**). Wakefulness and sleep were separated based on EMG power and on olfactory bulb gamma band power, which provides a movement-independent measure of arousal confirming the identification of sleep in head-fixed conditions. Both OB gamma and EMG were low during sleep (**Extended Data** Fig. 1a,b). REM (Rapid Eye Movement) sleep was identified based on high values of the ratio of theta-band to delta-band power in the hippocampus during sleep. During Non-REM (NREM) periods, we observed as expected a large increase of the slow wave and delta-band power compared to wakefulness and REM in the cortex, confirming the validity of our scoring approach (**Extended Data** Fig. 1c-e). Automated sleep scoring was performed in freely moving and head-fixed conditions in the same animals. After about two weeks of head fixation training of up to four hour sessions, we obtained very similar proportions of NREM sleep and wakefulness as in freely moving conditions (**Fig. 1j**), with sleep occurring towards the end of sessions (**Extended Data** Fig. 1a-c). In freely moving animals, NREM sleep bouts lasted up to a few minutes but these longer bouts were rare during head-fixation (**Extended Data** Fig. 1f). REM sleep was also observed under head-fixation but about twice less often than in freely moving mice (**Fig. 1j**). During two-photon imaging, mice were rarely awoken by sounds as only 2.00 ±0.98 % of sound presentations resulted in animals awakening and marginally affected sleep duration (**Extended Data** Fig. 1g,h). This allowed us to identify large epochs of sleep and wakefulness during both silence and sound presentation periods (**Fig. 1k**) during which we could record neural activity of the same population of neurons, visualized using Rastermap^17^ (**Fig. 1l**). As reported previously^13,18^, layer 2/3 auditory cortex spontaneous activity in the awake state was characterized by a mixture of desynchronized activity and population bursts (**Fig. 1l**), and we observed qualitatively the same structure in NREM sleep (**Fig. 1l**).

**Figure 1:**
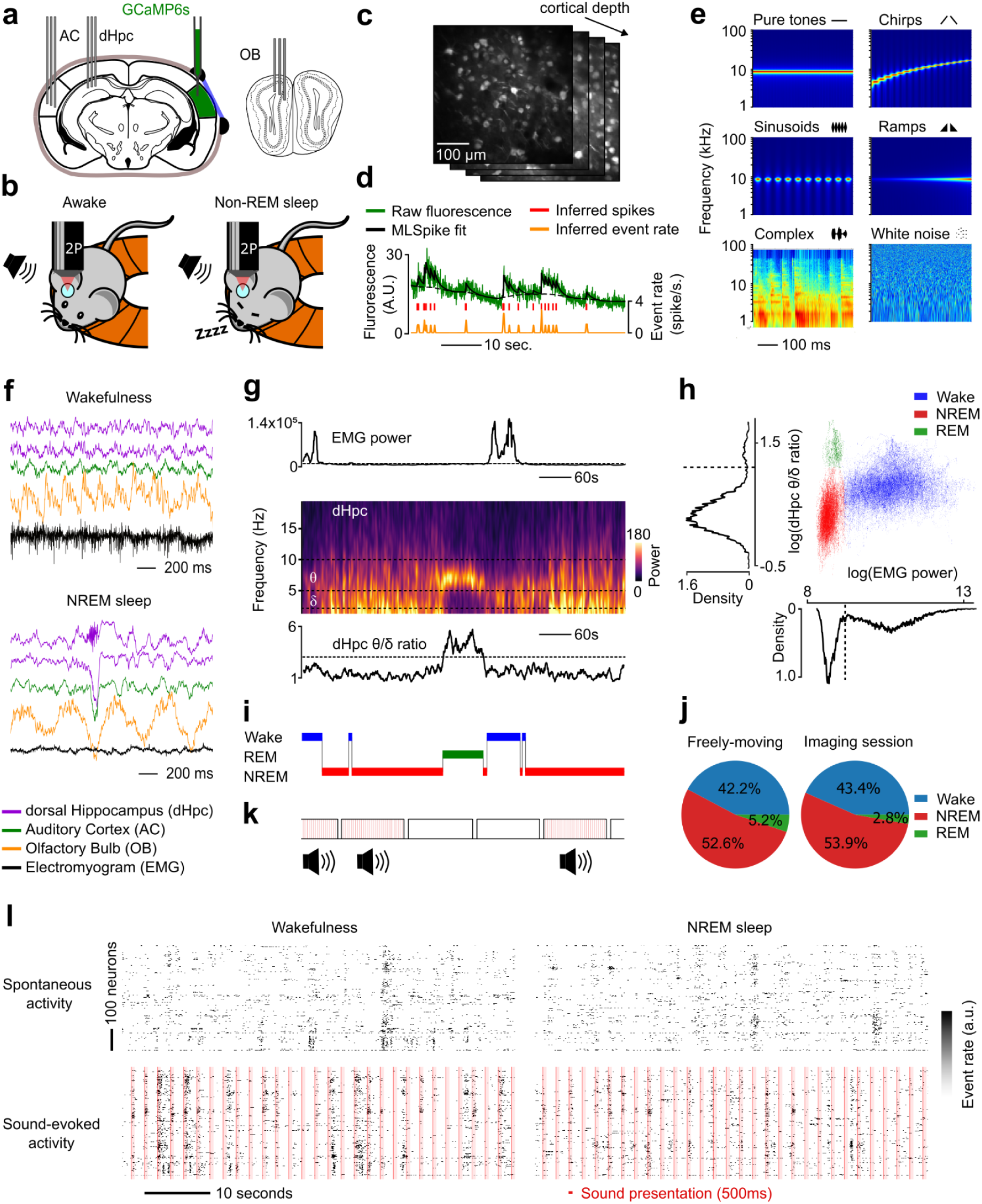
Robust sleep during head-fixed two-photon calcium imaging. **a,** Sites of Local Field Potential (LFP) electrodes implantation and AAV-GCaMP6s injection. A cover slip was placed over the right auditory cortex for imaging. AC = Auditory Cortex, dHpc = dorsal Hippocampus, OB = Olfactory Bulb. **b,** Recording setup. Mice were allowed to move on a treadmill while head-fixed under the objective of a two-photon microscope. Sounds were played when the animals were awake and asleep. **c,** Two-photon Ca^2+^ simultaneous recording of up to 1200 layer 2/3 neurons expressing GCaMP6s in 4 planes of different depth at 23 Hz. **d,** Deconvolution of calcium traces with MLSpike. Raw fluorescence signal is fitted with an exponential kernel to infer spike timings. Spike counts are binned in 43 ms windows and smoothed with a 100 ms Gaussian filter. **e,** Spectrograms of 6 example sound stimuli from different categories : pure tones, chirps, sinusoids, ramps, complex, and white noise. **f,** LFP example recordings at 1250 Hz of an animal in the awake and the NREM sleep state. **g.** The evolution of EMG power (top) reflects the arousal state of the animal. The spectrogram (middle) shows the evolution of theta and delta oscillation power changes in the dorsal Hippocampus with sleep stage changes, as the theta/delta ratio (bottom) increase reflects REM sleep. **h,** The bimodal distribution of EMG power identifies periods of sleep with low power and wakefulness with high power. The bimodal distribution of dHPC theta/delta ratio during sleep unveils periods of NREM (lower values), and REM sleep (higher values). The distribution of these variables outlines the 3 distinct arousal regimes. **i,** 10 min long extract of the hypnogram resulting from the sleep-scoring procedure. The animal falls into NREM sleep before entering REM sleep and wakes up. **j,** Proportion of time spent in each arousal state. Head-fixation did not change the time proportion spent asleep with regard to the freely-moving condition, but decreased by half the quantity of REM sleep. **k,** The recording was separated in 2 min long blocks where either sound stimuli were all presented once (red bars), or no sounds at all were presented (blank). The type of block was selected in a pseudo-random fashion and independently from the arousal state of the animal. **l,** Population raster plot of simultaneously recorded neurons during blank (top) and sounds (bottom) blocks, while the animal was awake (left) or asleep (right).

### Reduced sound discriminability during NREM sleep

We first compared sound responses in the auditory cortex across sleep and wakefulness. Given the small amount of REM sleep in imaging sessions, we focused the analysis on NREM sleep and excluded REM events. Plotting trial-averaged sound responses of specific neurons, we observed, as previously reported^6^, that neurons which responded to particular sounds in wakefulness respond to the same sounds in sleep **(Fig. 2a)**. These responses however had a weaker firing rate (**Fig. 2a**). In some, rarer cases weaker responses seen in wakefulness almost disappeared in sleep (**Fig. 2a**). Overall, the population firing rate was approximately halved (55.9%) across all sounds in sleep (**Fig. 2b, c**) indicating that despite qualitative preservation of sound responses, the neural activity carrying sound information is dampened in sleep. We estimated the sound-specific response ratio by dividing all sleep response amplitudes by the response ratio for the sound driving the largest response in wakefulness (**Fig. 2b**). We thereby observed a stronger reduction for simple, artificial sounds than for complex sounds (**Fig. 2b**). This reduced firing rate may result in a decrease of available sound information. To quantify this we used a cross-validated logistic regression classifier to measure the fraction of single trial population response for which the presented sound was correctly decoded. We observed a ∼15% decrease in decoder accuracy during NREM (**Fig. 2d**) thus indicating reduced sound information available in local neural populations of the auditory cortex. Misclassification, like the decrease of response gain (**Fig. 2b**), affected all sounds (**Extended Data** Fig. 2a-b), and was more likely to occur when population responses were weak (**Extended Data** Fig. 2e,f). As a point of comparison, we performed the same analysis on previously acquired data across wakefulness and isoflurane anesthesia^13^, which showed a massive sound information drop (> 3 fold) under anesthesia (**Fig. 2e**). Moreover, whereas sleep confusion matrices had a similar structure as in wakefulness, anesthesia generated systematic classification errors (**Extended Data** Fig. 2a,c). This shows that the reduction of sound responses observed in sleep results in a mild loss of sound information and is qualitatively different from the profound disruption observed under anesthesia.

**Figure 2:**
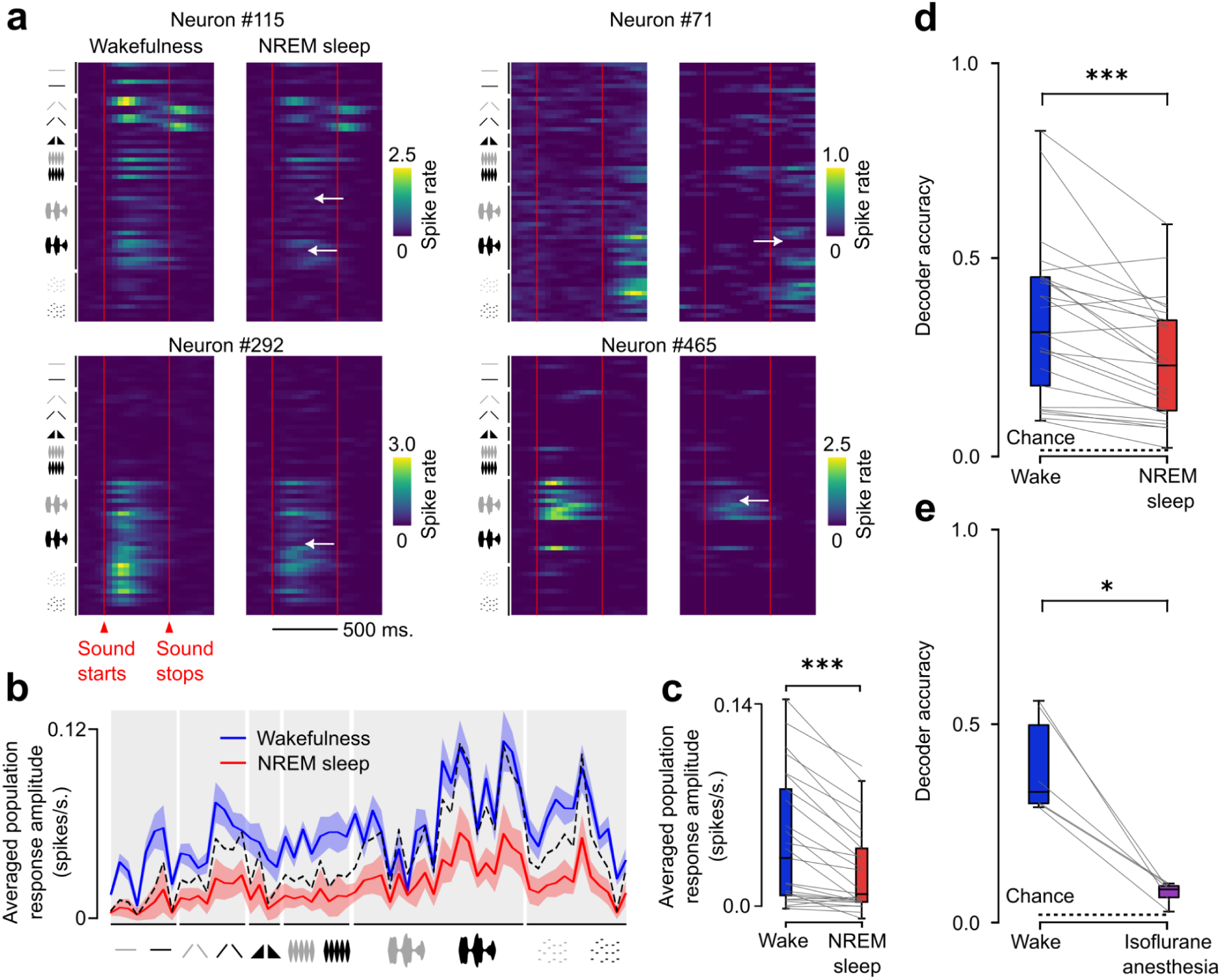
Conserved but reduced sound discriminability during NREM sleep. **a,** Averaged sound-evoked responses of 4 example neurons to each of the 60 stimuli presented (rows) sorted per category. Neurons respond to the same stimuli when the animal is awake or in NREM sleep, except for rare occasions where some responses almost disappear in NREM sleep (white arrows). **b,** Mean firing rate of neuronal population response is stronger in wakefulness than in NREM sleep for each of the 60 sounds presented. Solid line is the average over the 25 recording sessions, colored area is the standard error of the mean. Rescaling of the NREM curve so that its maximum corresponds to the maximum wake responses (dashed black line) highlights a stronger decrease of responses to all sound categories in NREM sleep relative to complex sounds. **c,** Average population firing rate over all sounds is stronger in wakefulness than in NREM sleep (Wilcoxon paired test, p=3.3×10^−6^, n=25 recordings). **d-e,** Linear support vector classifiers trained to decode sound identity from single trial neuronal population activity perform above chance level (dashed black line) but slightly better in wakefulness than in NREM sleep (d, Wilcoxon paired test, p=2.2×10^−5^, n=25 recordings), and accuracy drastically drops isoflurane anesthesia (e, Wilcoxon paired test, p=3.1×10^−2^, n=6 recordings).

### NREM sleep largely preserves the geometry of sound representations

Information loss can either be due to a modification of the geometry of representations (i.e. a recomposition of similarity relationships between sounds) or simply to a decrease of signal-to-noise ratio. In order to determine which of these two mechanisms underlies information filtering in sleep, we used a noise-corrected metric of similarity^19^, insensitive to the norm of activity pattern, to compare the geometry of sound representation across awake and NREM sleep states (**Fig. 3a**). The properties of the chosen metric allow identifying modifications of representation geometry itself independent of modifications of representation amplitude or variability, which relate the signal-to-noise ratio. Using this metric, we computed representation similarity analysis (RSA) matrices for the presented sounds, which summarize the representation geometry in the awake and NREM sleep states (**Fig. 3b**). We observed that RSA matrices were very similar across states (**Fig. 3b**), indicating that despite the small drop in information content, the structure of sound representation geometry in the awake auditory cortex is preserved in NREM sleep. We then computed the noise corrected similarity between sleep and wake sound representations to generate cross-state RSA matrices (**Fig. 3c**) and measured an average similarity level of 0.82 ± 0.09 for the same sounds across the two states (**Fig. 3d**). Thanks to our noise correction, the maximum similarity level of 1 would be reached with a confidence interval of 0.96 to 1.02 if representations are identical (**Fig. 3d**). Hence, while the differences in the representation of sounds in NREM sleep and wakefulness are subtle, they can be statistically detected. As a comparison, the representation similarity across wakefulness and isoflurane anesthesia, based on our previous data^13^, is much lower 0.33 ± 0.31 (**Fig. 3e**). Moreover, the structure of representation geometry is widely modified by anesthesia^13^ (**Extended Data** Fig. 3). Therefore, while anesthesia profoundly disrupts representation geometry, contributing to a large drop in sound information in the auditory cortex, NREM sleep preserves representation geometry and only marginally modifies sound response patterns themselves. This shows that despite the identified effects of NREM sleep on sound representations, they remain largely compatible with awake representations. Therefore a reorganisation of auditory cortex sound representations in sleep is not a mechanism that can explain auditory disconnection.

**Figure 3:**
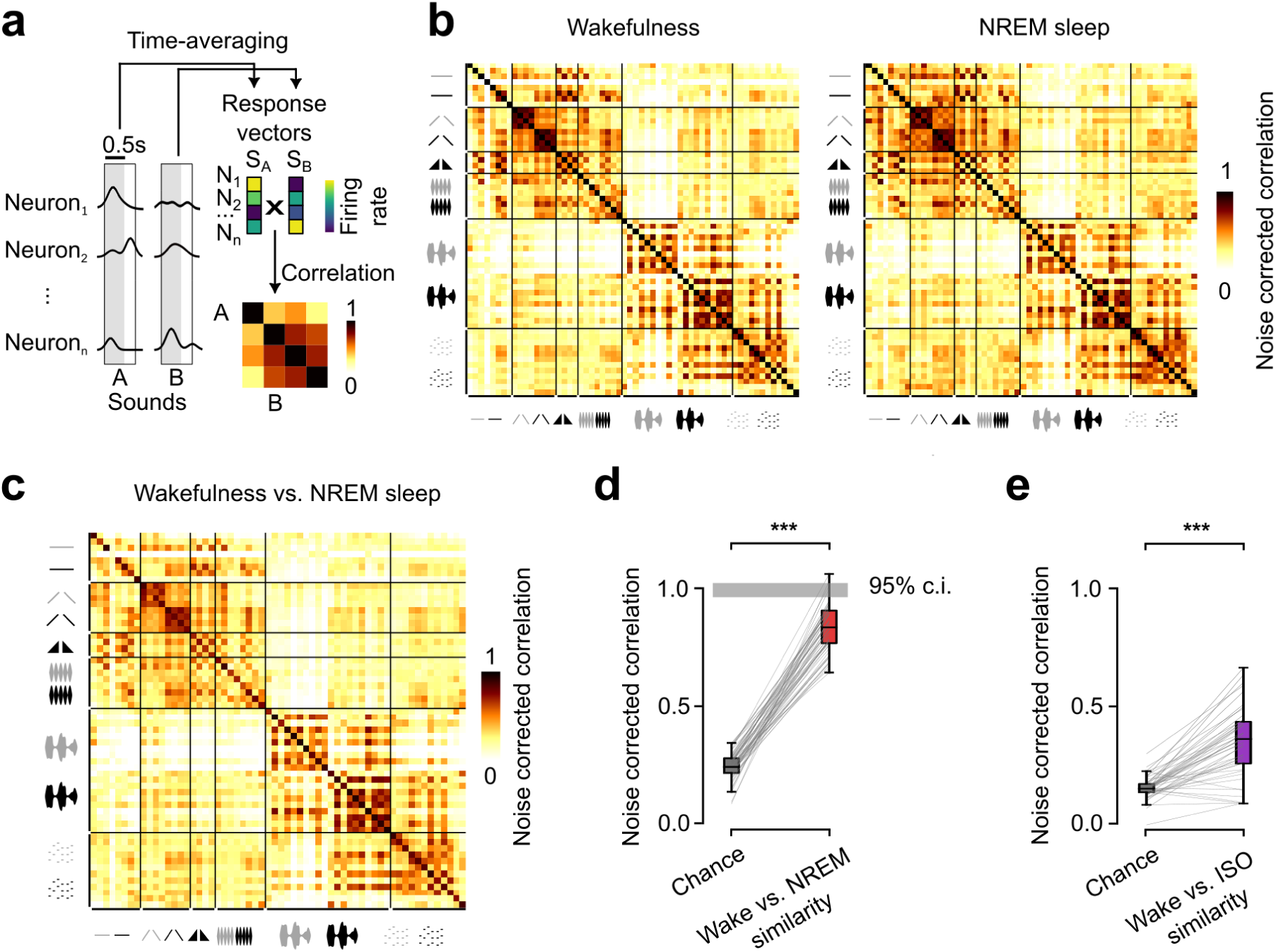
NREM sleep largely preserves the geometry of sound representations. **a,** Schematic explaining the construction of correlation matrices. Neuronal activity is averaged over time between sound presentation onset and 250ms after offset and concatenated into a vector for each sound. A high correlation value between two sound response vectors indicates a high similarity of neuronal population activation pattern. **b,** Noise corrected correlation matrices reveal identical similarity relationships between sounds in wakefulness (left) and NREM sleep (right). **c,** Matrix of pairwise noise corrected correlation between wakefulness and NREM sleep sound response vectors. **d,** Noise corrected correlation between same sound for wake and NREM sleep response vectors (diagonal of the matrix **c**) reveals a similarity level largely above chance (Wilcoxon paired test, p=1.6×10^−11^, n=60 sounds), but slightly below maximum expected level if representations are identical (95% confidence interval marked as grey area centered around 1). **e,** Noise corrected correlation between same sound wake and isoflurane induced anesthesia response vectors is slightly above chance (Wilcoxon paired test, p=3.1×10^−12^, n=50 sounds).

### Spontaneous and evoked activity cover different subspaces in NREM sleep

It was recently observed that, contrary to the wake state in which spontaneous and sound-evoked activity are clearly distinct^20^, under isoflurane and ketamine anesthesia, they produce very similar neural population patterns^13^. This phenomenon could explain sensory disconnection because if evoked activity is indiscriminable from on-going activity in the auditory cortex, then it is not possible for downstream target areas to identify if a sound is present or not. To determine if a similar phenomenon occurs in NREM sleep, we collected spontaneous activity from extended periods of silence, and sound-evoked activity patterns systematically in time bins of 43ms (imaging frame duration) and decomposed them using principal component analysis, separately for wakefulness and NREM sleep. Projections of spontaneous and sound-evoked patterns in the reduced space of the second and third components qualitatively showed that spontaneous and evoked activity patterns are distributed in different regions of the activity state space both in wakefulness and NREM sleep but not under isoflurane anesthesia (**Fig. 4a**). This suggests that spontaneous and evoked activity in sleep have little overlap, as in wakefulness. To quantify this further, we measured the cross-validated amount of sound-evoked activity variance captured by the spontaneous activity subspace (**Fig. 4b**). The spontaneous subspace was defined as the activity captured by the N first principal components which accounted for more variance than expected by chance (**Extended Data** Fig. 4a, chance is estimated based on a shuffling of cells activity in time in spontaneous activity patterns, see methods). This approach allows us to estimate the dimensionality of the spontaneous activity subspace in the recorded population, and the extent to which evoked activity is included into the spontaneous activity subspace (**Fig. 4b**). The reasoning is that if spontaneous and evoked activity are fully overlapping, the amount of variance captured by the projection of evoked activity onto the spontaneous activity subspace should be equal to the amount of variance captured by the projection of spontaneous activity onto its own subspace. Dimensionality of spontaneous activity was variable across sessions due to variations of neural population size but was large (several 10’s to a few 100’s of principal components) both in the awake and NREM sleep states (e.g. **Fig. 4c**, **Extended Data** Fig. 4a,b), while under isoflurane anesthesia, at maximum, the first ∼20 components accounted for most of the significant variance in the spontaneous activity data (**Fig. 4c, Extended Data Fig. 4a,b**). In wakefulness, as expected from previous observations in the auditory^13^ and visual^20^ cortex, the projection of evoked activity onto the spontaneous activity subspace captured only a small fraction of evoked activity (**Fig. 4c,d**), and only the very first principal components of spontaneous activity significantly captured evoked activity (**Fig. 4c**). Strikingly, the same analysis in NREM sleep demonstrated also a limited overlap between spontaneous and evoked activity (**Fig. 4c,d**), contrary to anesthesia (**Fig. 4c-e**). Notably, spontaneous activity in wakefulness and sleep populated clearly different subspaces (**Extended Data** Fig. 4c) in line with the profound change of cortical dynamics in NREM sleep compared to wakefulness, while sound-evoked activity in NREM and wakefulness shared their subspaces to a greater extent (**Extended Data** Fig. 4d,e) as expected from the similarity of population response patterns across these states. Overall this shows that no gating mechanism based on the similarity of spontaneous and evoked activity is implemented in NREM sleep.

**Figure 4:**
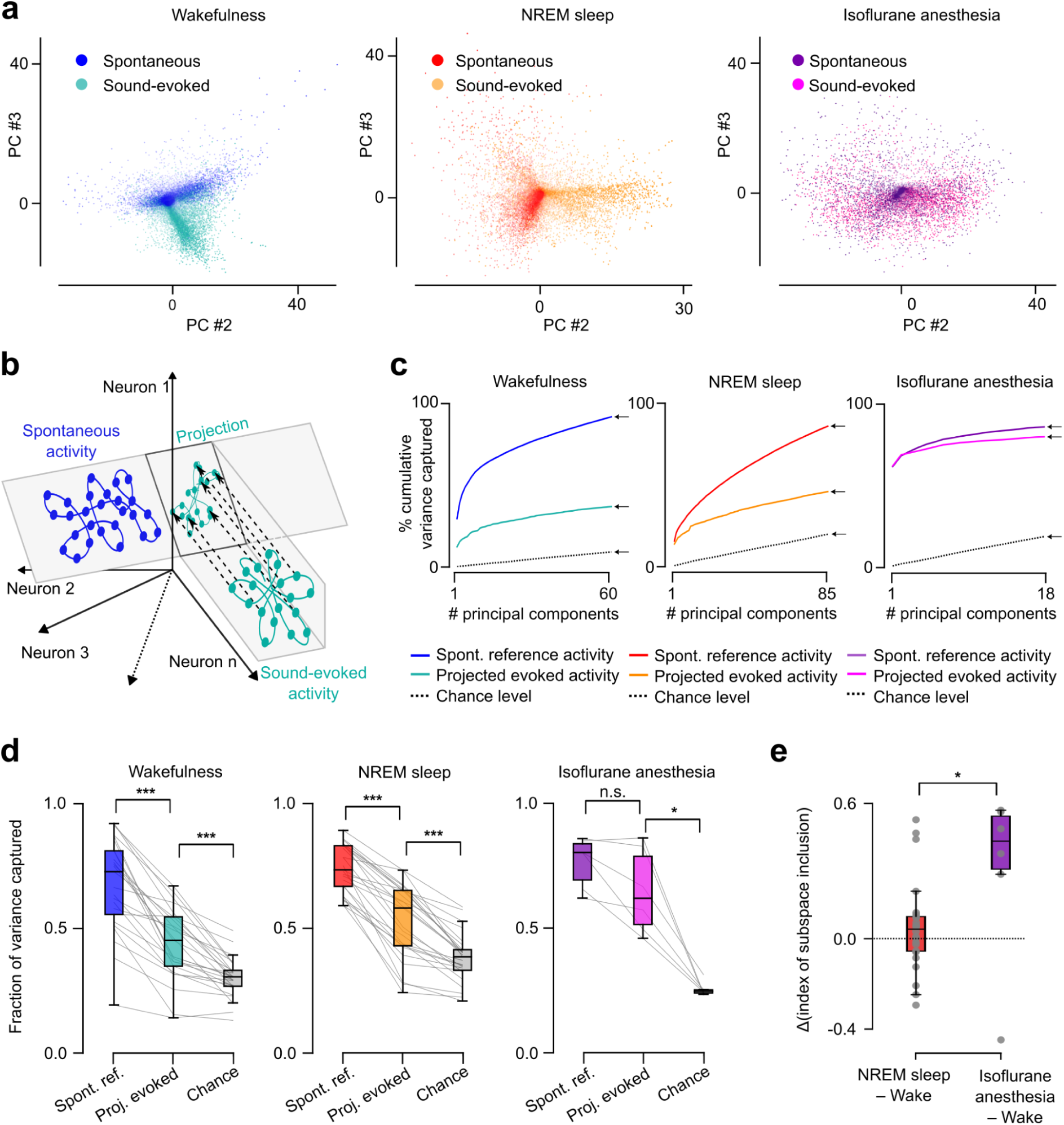
Spontaneous and evoked activity cover different subspaces in NREM. **a,** 2D example projection of neuronal activity in principal component (PC) space reveals distinct dimensional occupancy for spontaneous and sound-evoked activity in wakefulness (left) and NREM sleep (middle), but overlap in isoflurane induced anesthesia (right). **b,** Schematics explaining the calculation of the fraction of sound-evoked variance captured by the spontaneous activity space. The sound-evoked activity is projected into a subspace of the spontaneous activity space defined as the ensemble of significant principal components derived from a first subset of spontaneous activity data. To estimate the level of overlap of spontaneous and sound-evoked activity, we compared the variance of this projection with the variance of the projection of a second subset of spontaneous activity data that serves as a control for the maximum possible overlap. **c,** The fraction of variance captured from the sound-evoked activity projected on the PCs of spontaneous activity is much lower than the maximum overlap expected in both wakefulness (left) and NREM sleep (middle) and this gap increases when adding successive PCs. Under isoflurane anesthesia (right) even the last PCs show a strong overlap between spontaneous and evoked activity. Examples from single recordings. The arrows point towards the values of variance captured by the entirety of the defined reference spontaneous subspace. **d,** Quantification of fraction of variance captured from all recordings in the different states (Wilcoxon paired test, Wakefulness : “spontaneous reference” to “Projected evoked” : p=6.0×10^−8^, “Projected evoked” to “chance level” : p=1.2×10^−7^, n=25; NREM sleep: “spontaneous reference” to “Projected evoked” : p=6.0×10^−8^, “Projected evoked” to “chance level” : p=6.0×10^−8^, n=25 recordings; Isoflurane induced anesthesia : “spontaneous reference” to “Projected evoked” : p=6.2×10^−2^, “Projected evoked” to “chance level” : p=3.1×10^−2^, n=6 recordings). **e,** The difference in sound-evoked subspace into spontaneous subspace inclusion level between wakefulness and NREM sleep is smaller than between wakefulness and isoflurane anesthesia (Mann-Whitney rank test, p=3.3×10^−2^). Positive values indicate increased inclusion of sound-evoked activity in spontaneous subspace compared to wakefulness.

### Intermittent gating of local population responses during NREM sleep

Our results so far indicate that auditory cortex sound responses have preserved geometry during sleep and are not masked by spontaneous activity. Thus, the auditory cortex processes sound information with little distortion, apart from a reduction of sound-related information due to reduced neuronal response amplitudes. In single neurons, this reduction of response amplitude in sleep reflects both a decrease of single trial response probability (i.e. whether or not a neuron responds) and a decrease of the single-trial response amplitude (i.e. how strongly a neuron responds, **Fig. 5a,b**). However, from a neural population perspective, response variations could happen independently in each neuron or could be coordinated, leading, in the first case, to a mild reduction of the signal-to-noise ratio, and in the second case to potentially severe disruptions of information transmission at specific times when all neural responses are globally reduced. To qualitatively evaluate these two alternatives, we plotted single-trial population responses, sorted according to the norm of the population response vector (**Fig. 5c**). These plots clearly indicated that population responses are more robust in wakefulness than in sleep (**Fig. 5c**), and that fluctuations of population response amplitude are much larger in NREM sleep, with some sound presentation leading to strong population responses, comparable to wake, and others to an apparent absence of response (**Fig. 5c**). To quantify this more precisely, we selected, for the ten most responsive sounds, the 30% trials with the largest population response amplitude, and averaged them to build a template response pattern, representative of well-driven responses. We then measured the cosine distance between every single trial pattern and the template response pattern to estimate the amount of sound-related information conserved at the single trial level. While in wakefulness most trials produced response patterns very similar to the template, in sleep this similarity dropped for a large fraction of trials. The drop in similarity was larger during sleep than expected if single-neuron trial-to-trial variability was uncoordinated as evaluated with independent trial shuffling across neurons (e.g. **Fig. 5d**). Cosine similarity was tightly correlated to the baseline-subtracted population firing rate (**Extended Data** Fig. 5a-c) indicating that low cosine similarity corresponds to low responsiveness of the population above spontaneous baseline firing. Moreover, as we observed qualitatively on activity heatmaps (e.g. **Fig. 5c**) a number of single trial responses had a cosine similarity with the template response pattern close to 0 (e.g. **Fig. 5d**) indicating that the corresponding activity pattern is close to orthogonal to the main response axis. Together, these demonstrate that coordinated fluctuations in response strength are larger during sleep, leading to a complete lack of response for certain sound presentations.

**Figure 5:**
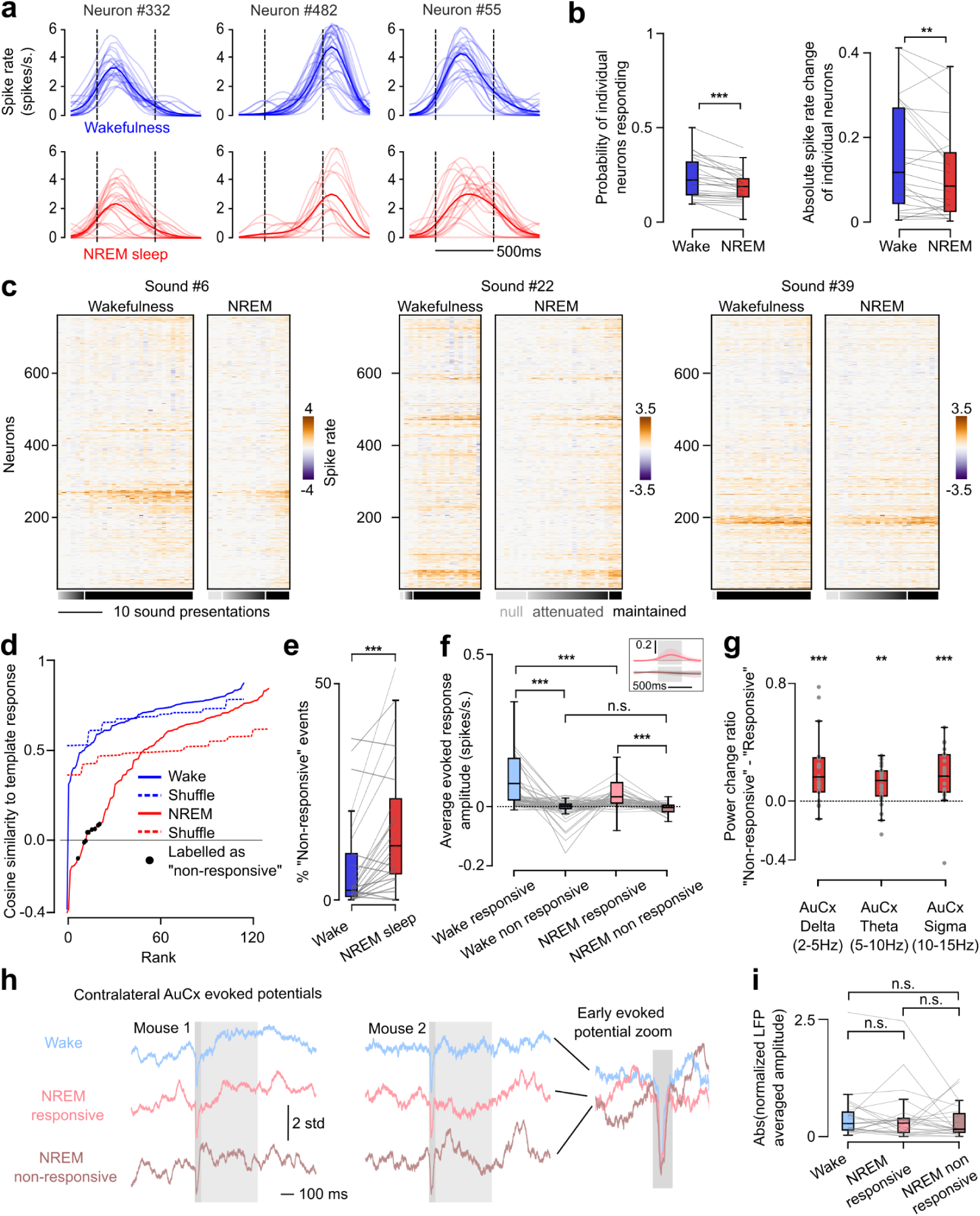
Intermittent gating of population responses during NREM sleep. **a,** Example of neuronal responses to sounds highlighting the reduced evoked response amplitude in NREM sleep. Thin traces are single trial sound presentations, thick line is the average over trials. **b,** Average probability of neurons having a non-zero response to sounds is higher in wakefulness than in NREM sleep (left, Wilcoxon paired test, p=5.4×10^−5^, n=25 recordings). Average response amplitude when non-zero is also higher in wakefulness than in NREM sleep (right, Wilcoxon paired test, p=6.7×10^−3^, n=25 recordings). **c,** Single trial neuron population responses relative to baseline for example sounds reveal events of coordinated reduction of sound-evoked activity in NREM sleep that are not seen in wakefulness. Trials are sorted by increasing population response intensity. The greyscale bars qualitatively indicate the level of conservation of population responses from completely maintained to absence of response (null). **d,** Cumulative distribution of similarity between single trial population responses and the corresponding template sound–evoked response in NREM sleep shows more reduced similarity than in wakefulness. Values not significantly different from zero (indicating that the response is essentially orthogonal to the template) are labelled as “non-responsive” and indicate suppression of sound encoding at the population level (black dots). Dashed lines indicate the similarity values distribution when neuron response intensities are independently shuffled between trials. **e,** The proportion of trials for which the population is classified as “non-responsive” is increased in NREM sleep compared to wakefulness (Wilcoxon paired test, p=1.2×10^−4^, n=25 recordings). **f,** Population response amplitude drops during “non-responsive” events (Wilcoxon paired test, wakefulness : p=1.4×10^−10^, n=82; NREM sleep : p=6.5×10^−23^, n=154) and is as low in wake and NREM sleep (Wilcoxon paired test, p=8.2×10^−1^, n=69). Amplitude remains higher in wakefulness than NREM sleep when the population is responsive (Wilcoxon, paired test, p=2.1×10^−27^, n=230). Inset shows the average sound-evoked response during “responsive” (top) and “non-responsive” (bottom) events in NREM sleep. **g,** Delta, Theta, and Sigma oscillation power in the contralateral AuCx in a 1.5s long window centered around sound presentation increase in “non-responsive” compared to “responsive” events (Wilcoxon paired test, Delta : p=7.3×10^−5^ ; Theta : p=1.3×10^−3^, Sigma : p=1.3×10^−4^, n=25 recordings). **h,** Averaged LFP around sound presentations for 2 example animals recorded in the contralateral auditory cortex during wakefulness, and “responsive” and “non-responsive” events in NREM sleep. Raw LFP traces were divided by their averaged value over the entire recording to allow comparison between conditions and recordings. Light grey area shows sound presentation time, dark grey area is the 50ms interval after sound onset where a peak in the sound-evoked potential occurs. **i,** Mean amplitude of the early sound-evoked potential in the 50ms window is comparable between the three conditions (Wilcoxon paired test, “wake” to “NREM responsive : p=7.1×10^−1^; “wake” to “NREM non-responsive” : p=1.6×10^−1^; “NREM responsive” to “NREM non-responsive” : p=3.3×10^−1^, n=25 recordings).

To identify these non-responsive events under statistical control, we used a bootstrap test to evaluate trials with response patterns orthogonal to the direction of well-driven, template sound response patterns (**Fig. 5-d**, see methods). This showed that 17 ± 14 % of sound presentations in NREM sleep resulted in an absence of response while this fraction was much smaller in wakefulness (7.8 ± 11 %, **Fig. 5e**) and often close to zero in well-driven wake populations (**Fig. 5c-e**). Trials with these null responses (i.e. orthogonal to the main axis of response) can be seen as non-responsive trials because their baseline-corrected population activity is close to 0 (**Fig. 5f**). This indicates no changes relative to ongoing, spontaneous activity and thus a lack of information about the sound. We observed that non-responsive trials contribute to a fraction of the sound information loss in the auditory cortex during sleep, the remaining fraction being imputable to the other trials with a reduced, but non-zero response gain compared to wakefulness (**Extended Data** Fig. 6a).

Finally, we asked if the lack of neural response to sounds we observed in these local neural populations is related to indicators of local or global state in the brain and body. Non-responsive trials were associated with a weak but significant increase in baseline spontaneous activity amplitude in the imaged neural population just before the sound (**Extended Data** Fig. 6b-d), that was not associated with a particular change of the spontaneous activity dynamics as estimated from population activity projections onto its first principal components (**Extended Data** Fig. 6c). At a more global scale, power in low-frequency bands, including the delta (2-5 Hz) and the sigma (10-15 Hz, spindle associated) bands was increased in the contralateral auditory cortex during non-responsive trials in NREM sleep (**Fig. 5g, S5d,e, Extended Data Fig. 6e,f**). These indicate that full blockade of sound information observed locally is more probable during oscillatory patterns typically associated with deeper sleep and sensory disconnection^21,22^. Non-responsive trials were not simultaneous with spindle events (**Extended Data Fig. 7**) and so are not directly due to the spindles themselves, corroborating previous observations that spindles do not affect sound-driven activity in the auditory cortex^7^. Non-responsive trials also did not necessarily co-occur with delta waves in the contralateral auditory cortex (**Extended Data Fig. 8**). We moreover found that the early auditory evoked potential in the contralateral auditory cortex had identical amplitudes whether or not an ipsilateral disconnection was observed in the imaged population (**Fig. 5h,i**). Finally, slight muscle contractions elicited by the sounds were still present during non-responsive trials (**Extended Data Fig. 6h**). These observations demonstrate that some auditory signals are preserved at least up to the thalamic input to the cortex during non-responsive trials and therefore, that the intermittent absence of response observed locally in the auditory cortex does not correspond to a global disconnection of the thalamo-cortical axis from auditory inputs.

## Discussion

In this study, we reveal that sounds are encoded with a highly similar geometrical structure between wake and sleep but with a small but significant drop in accuracy linked to an overall reduction in firing rate (**Fig. 2**). We show that this loss is explained by a mixture of reduced overall activity and coordinated silencing within the neuronal population (**Fig. 5**). This silencing was simultaneous with an increase in broadband low-frequency brain oscillations power especially delta and sigma rhythms (**Fig. 5**).

Previous work has shown that auditory cortex neurons preserve their sound responses during sleep^5,6^. We extend this finding to show at the population level that the geometry of auditory representations is also conserved between sleep and wake. First, the relations between sounds, as quantified with Representation Similarity Analysis(**Fig. 3**), is conserved in spite of differences in spontaneous activity and ongoing dynamics. Moreover, spontaneous and evoked activity are well separated during sleep, excluding the possibility that sensory disconnection in sleep results from a masking of evoked responses by spontaneous activity (**Fig. 4**). This result extends to the NREM sleep state the recent observation of a clear difference between evoked and spontaneous activity in awake animals^20^, and strongly contrasts with anesthesia during which evoked activity strongly resembles spontaneous activity, both in the auditory^13,23^ and visual^24^ cortex. These observations are consistent with the profound anesthesia-induced disruptions of auditory representations found even at the earliest stages of auditory processing^25^ and show that modulations of sensory processing in sleep and under anesthesia are mechanistically distinct.

Strikingly, we observed that the preserved geometry of sound representation in layer 2/3 neurons of the auditory cortex is paralleled by an approximately 50% reduction in firing in response to sounds. This reduction of the neural signal strength leads to a reduction in the accuracy with which sounds can be decoded based on neural population activity during sleep. Quantitatively, the sleep-induced reduction in cortical responses to sounds that we report is large compared to previous reports of no change^5,6^, or changes restricted to soft sounds^26^ or to late phases of responses^27,28^. These studies were performed with electrophysiology which tends to sample deeper layers whereas our recordings were restricted to layer 2/3. This may account for the difference since a previous electrophysiological study carefully sampling across layers identified intermittent shutdowns of layer 2/3 while layer 5 remained active^29^. Moreover, whereas electrophysiology shows preserved onset responses and reduced late responses^27,28,30^, the slow temporal resolution of calcium imaging pools these periods together, thus emphasizing the net reduction in responsiveness.

Despite the reduced response amplitude, the preserved geometry of sound representations during NREM sleep clearly shows that the auditory cortex continues surveilling the environment based on a processing of acoustic information similar to processing performed wakefulness. This is consistent with the demonstration in human subjects that even the extraction of sophisticated auditory features such as word identity continues to operate during sleep^10^ all the way up to nonconscious preparation of motor responses^31^. This mechanism could be sense-specific, as the ear remains physically receptive during sleep and audition allows to monitor distal events. This could confer strong evolutionary advantages since awakening to meaningful stimuli^8,9^ mitigates the risks associated with sensory disconnection during sleep. However, isolation of the brain from the environment is thought to be crucial to enable processing that faithfully reflects neural internal constraints such as homeostatic plasticity, memory consolidation or emotional regulation^12,32^. In the auditory cortex, the intermittent coordinated reduction in responsiveness that our study uncovers could be a way for this cortical region to recuperate without needing to be permanently switched off during NREM sleep.

Importantly since spontaneous activity is largely orthogonal to evoked responses during sleep (**Fig. 4**), the interferences of sound responses with the internally-generated sleep-dependent processes such as memory consolidation^33,34^ may be minimized. Our data does not precisely delineate the scale at which the intermittent gating of sensory information occurs in the cortex. We show that at least contralateral thalamic inputs and early cortical potentials are preserved during ipsilaterally observed gating events (**Fig. 5**) indicating that these events are not a full shutdown of auditory processing, and are rather restricted to the cortex. We found that local population gating events are correlated with contralateral oscillatory activity modulations (**Fig. 5g, Extended Data** Figs. 5d and 6e**,f**), albeit with low predictive power (**Extended Data** Fig. 6d). This suggests that auditory population responsiveness could be impacted by slow changes in vigilance state^35,36^ instead of specific time-locked events. On the other hand, since sleep can be accompanied by localized changes in the neural circuit^37^, our data is also compatible with the hypotheses that local patches of cortex intermittently go offline while others preserve monitoring. Independent of the scale at which this mechanism occurs, it provides a powerful way to combine the need for surveillance and the need for offline processing during sleep.

Our results raise the question of where sensory gating is happening since the weak reduction in responses with preserved population geometry that we observe seem insufficient to explain the sensory disconnection that is the hallmark of sleep. Our results support the hypothesis of a downstream, frontal gating^1,2^. Perturbations, whether sensory or artificial using TMS or optogenetics, show that frontal regions typically respond to sensory inputs with an evoked downstate that shuts off inter-area communication^38–40^. Therefore, sound responses in the auditory cortex may not be passed downstream in the processing hierarchy.

## Material and Methods

### Animals

Seven male C57BL/6 mice were used for experiments, all aged between 8 to 24-week-old, and had not undergone any other procedure before. Animals were housed in groups of two to four in controlled temperature and humidity conditions (21-23°C, 45-55% humidity), were maintained on a 12h light-dark cycle, and had *ad libitum* access to food, water, and enrichment (wooden logs, cotton bedding, and running wheel). All experiments were conducted during the light phase. All procedures were carried out in accordance with the French Ethical Committees #89 (authorization APAFIS#27040-2020090316536717 v1).

### Surgery

Mice were injected with buprenorphine (Vétergesic, 0,05-0,1 mg/kg) 30 min prior to surgery. Surgical procedures were carried out using 3.5% isoflurane for induction delivered in an airtight box, and were maintained at 1-2% isoflurane via a mask for the rest of the procedure. After induction, mice were kept on a thermal blanket during the whole procedure and their eyes were protected with Ocrygel (TVM Lab). Lidocaine was injected under the skin of the skull 5 minutes prior to incision.

For local field potential recordings, coated tungsten (.002” bare, .0040” coated) wires were chronically implanted intracranially in the olfactory bulb OB (centered on Bregma; A/P: 4.2mm, M/L: −0.5mm, D/V: 2.25mm), the dorsal hippocampus Hpc (centered on Bregma; A/P: −2.2mm, M/L: −1.7mm, D/V: 1.3mm), the primary auditory cortex AC (centered on Bregma; A/P: −2.5mm, M/L: −3.75mm, D/V: 1.25mm) with a motorized stereotactic arm (StereoDrive, Neurostar, Tubingen, Germany), and at the surface of the cerebellum for ground and references. Silver wires (.005” bare, .0070” coated) were wrapped around the dorsal neck muscles for electromyography. All electrodes wires were pinned to 16 channels EIB (Omnetics, Minneapolis, MN) fixated on the skull of the animal with dental cement (Super-Bond, Sun Medical, Japan).

For calcium imaging, a 5 mm diameter circular craniotomy was performed above the auditory cortex. Three to six injections of 150nL of AAV1.Syn.GCaMP6s.WPRE (Vector Core, Philadelphia, PA; 10^13^ viral particles per ml; used diluted 30x) were done at 30 nl/min with pulled glass pipettes at a depth of 500µm and spaced every 500µm to cover the large surface of the auditory cortex. The craniotomy was sealed with a circular glass coverslip. The coverslip and head-post for head-fixation were fixed to the skull using cyanoacrylate glue (3M Vetbond^TM^) and dental cement (Super-Bond, Sun Medical, Japan). After surgery, mice received a 30% glucose solution, and metacam (1 mg/kg) through subcutaneous injection. Mice had metacam delivered via their drinking bottle in their home cage for five days starting the day following the surgery. Mice were allowed to recover for a week without any manipulation.

### Two-photon imaging

Calcium imaging was performed using a custom two-photon microscope (AODScope, Karthala Systems, Paris, France) equipped with an acousto-optic-deflector scanner combined with a pulsed laser (MaiTai-DS, SpectraPhysics, Santa Clara, CA) set at 920 nm. We used a 16x Nikon CFI LWD water dipping objective (MRP07220), which provided a field of view up to 450×450µm. Images were acquired either at 30Hz in monoplane mode, or at 23Hz in multiplane mode with 50µm depth difference between planes. Laser power reaching the sample was between 80 and 120mW.

### Local Field Potential recordings

Local field potentials were recorded through a FPGA acquisition board from OpenEphys connected to a RHD 16-channel recording headstage from Intan plugged into the EIB-16 embedded on the head of the animal. Signals were high-pass filtered at 0.1 Hz and recorded at 1250 Hz.

### Sleep-scoring procedure

Animal vigilance state was classified by a two-step automatic algorithm described and previously validated in Bagur et al., 2018^16^. In the first step, the high-frequency band (50-300 Hz) power in the EMG or the gamma band (50-70 Hz) power in OB oscillations was used to distinguish wakefulness from sleep. The instantaneous power of the corresponding frequency-band was computed as the Hilbert transform of the filtered signal, and smoothed using a 3 s sliding window. The distribution of these values were fit to a mixture of two Gaussian distributions normalized to each having an area of one, so that the intersection of these two distributions is used as a threshold to separate wakefulness and sleep independently from sleep to wake quantity ratio. Values below the threshold were classified as sleep while values above were classified as wakefulness. In a second step, LFP signal from the Hpc during sleep periods only was filtered in the Delta band (2-5 Hz) and the Theta band (5-10 Hz) and their respective instantaneous power was computed as their Hilbert transform and smoothed using a 2s sliding window. A single normal distribution was fit to the distribution of the ratio of Theta / Delta power. The NREM/REM threshold was placed at the point where the fit explained less than 50% of the data, values below the threshold were classified as NREM sleep while values above were classified as REM sleep. Periods of NREM and REM sleep shorter than 3s were merged into the surrounding periods to avoid artificially short epochs.

### Experimental protocol and auditory stimulation

After the post-surgery recovery period, mice had their LFP signals recorded in freely-moving conditions in a 35 x 18 cm home-cage with litter and food pellets, for 6 to 8 h during two consecutive days to quantify sleep quantity. Then, mice were habituated to head-fixation for progressively increasing durations (from 20 min to 3 h) over two weeks, either in a box or on a wheel. The box setup consisted of a 10 x 10 cm box with 5 cm high walls 3D printed with polylactic acid, its floor was covered with litter to absorb urine and feces. The wheel setup consisted of a 15 cm diameter toy horizontal running wheel to allow free locomotion of the animal, and was also covered with litter to absorb urine and feces. Each mouse was habituated to a single type of setup only. LFP signals were recorded during habituation sessions.

When an animal was sufficiently habituated to head-fixation to fall asleep and display REM sleep, it was used for imaging sessions. During the sessions, the mouse was head-fixed on the setup it was habituated to. The activity of neurons in the primary auditory cortex was recorded with two-photon calcium imaging through the cranial window, and LFP signals were recorded to monitor the vigilance state of the animal. The recording was split into 80 to 100 blocks of two minutes with either silence or sounds. Equal numbers of silence and sounds stimulation blocks alternated randomly to sample spontaneous and sound-evoked activity during wake and sleep throughout the session. Within a stimulation block, all sound stimuli were presented once in a randomized order. Each sound was 450 to 500 ms long, and sound onsets were separated by a 2 s interval within the block. Sounds were delivered at 192 kHz with a NI-PCI-6221 card (National Instrument) driven by a custom Matlab software through an amplifier and high-frequency loudspeakers (SA1 and MF1-S, Tucker-Davis Technologies). Sounds were calibrated in intensity at the location of the mouse ear using a probe microphone (Bruel & Kjaer).

The sound stimuli consisted of a collection of 60 predefined sounds divided into six groups (**Fig. 1e**): four pure tones (6, 8, 12, and 25 kHz), four frequency-modulated sounds (4–16, 4–25 kHz, upward and downward modulations), four ramping amplitude modulation sounds (8 and 12 kHz), four sinusoidal amplitude modulation sounds (sinusoidal modulation at 20 and 80 Hz, for two carrier frequencies of 8 kHz and 12 kHz), ten complex sounds, and white noises and derivatives (raw white noise, ramping up and down, filtered in the 6.4 to 19 kHZ and 8.6 to 12 kHz band, filtered around 12 and 25 kHz and ramping up and down, and filtered in the 13 to 24 kHz band with amplitude sinusoidal modulation at 40 Hz). All sounds (except for sinusoidal and ramping sounds) were played at “low” intensity (50 or 60 dB SPL) and “high” intensity (70 or 80 dB SPL).

### Data analysis

#### Calcium signal processing

Data analysis was performed using Python and MATLAB scripts. Artifacts related to horizontal movements were corrected frame by frame using rigid and non-rigid body registration. ROI were identified by applying Singular Value Decomposition (SVD) on the movie and finding peaks in the components, and were then cured manually. Registration and ROI identification were performed using the Suite2P algorithm^14^. Neuropil contamination was subtracted by applying the following equation : 𝐹_𝑐𝑜𝑟𝑟𝑒𝑐𝑡𝑒𝑑_(𝑡) = 𝐹_𝑚𝑒𝑎𝑠𝑢𝑟𝑒𝑑_(𝑡) − 0. 7𝐹_𝑛𝑒𝑢𝑟𝑜𝑝𝑖𝑙_(𝑡) to each ROI, where 𝐹 is the fluorescence and 𝑡 is time. These corrected traces were deconvolved using the MLSpike algorithm^15^ to estimate the putative timing of spikes trains underlying the recorded fluorescence. A time constant of calcium transient τ = 2s, and polynomial coefficients [p_2_,p_3_] = [0.73, −0.05] have been chosen in accordance with the published estimations for GCaMP6s dynamics^15^. Unitary spike coefficient a = 0.1 and coefficient adjusting for baseline drift compensation (0.2) were estimated to best fit the descending slope of experimental calcium spikes as well as the fluctuation of the baseline fluorescence. The number of spikes was then counted for each neuron within time bins corresponding to frames acquired by the microscope (33ms for monoplane acquisition, 43ms for multiplane acquisition), and this spike count was smoothed by convolution with a Gaussian kernel (std = 100ms, if not stated otherwise).

#### Stimulus decoding classifiers

Linear Support Vector Classifiers (LinearSVC) were trained to decode stimulus from neural activity. For each recording session, classifiers were fed with the population responses vectors to sound stimuli. Population response vectors were computed by concatenating the response of every neuron recorded in the session to each sound presented during the experiment. Responses were averaged in time over stimulus presentation and the following 250ms to take into account responses to sound offset. For each session, classifiers were trained with a stratified 5-fold cross-validation procedure : the dataset was randomly split in 5 non overlapping groups with the same number of stimuli and comparable proportion of each stimulus type. The classifier was trained on 4 of the groups and accuracy was tested on the remaining group. The procedure was repeated so that each group plays the role of the test set once. Accuracy of the classifier was computed as the weighted F1 score across stimuli in the test sets. The number of sound presentations were equalized between wake and NREM sleep conditions to ensure that observed differences in performance of the classifiers would not depend on the size of the training dataset between the two conditions.

#### Representation Similarity Analysis

To assess the level of similarity between neuronal patterns of responses to different sound stimuli, we implemented an estimation of the Pearson correlation coefficient correcting for trial-to-trial variability. The level of similarity between two neuronal patterns was computed as : 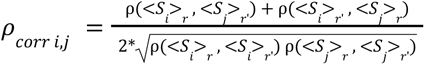, where 𝜌 denotes the Pearson correlation coefficient, *<S_*i*_>*_*r*_ is the population vector in response to sound *i* averaged over trials from the ensemble *r*. The ensembles *r* and *r’* randomly split the dataset in two halves with equal number of occurrences of each stimulus type. This metric allows to normalize the correlation level to the maximum correlation level that can be expected given the noise in single trial responses. Because responses during wake and NREM sleep do not share the same ensemble of trials, the similarity level of the responses between the two states can be written as : 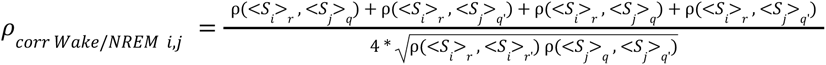, where *r* and *r’* split in half sound presentations occurring during wakefulness periods, and *q* and *q’* during NREM sleep periods. Similarity of coding was assessed when *i = j*, and its corresponding chance level was calculated by shuffling sound identity so that it is mismatched between the two states. The expected value obtained when representations are similar is computed by splitting wakefulness data in 4 non-overlapping groups of sound presentations *r*, *r’*, *q*, *q’*, and should be centered around 1.

#### Dimensionality of neuronal activity subspaces

To assess the dimensionality of neuronal activity subspaces we performed PCA on the time series of simultaneously recorded neurons during a specific condition (spontaneous activity recorded during blocks of silence, sound-evoked activity during sound presentations time, in wakefulness, NREM sleep, and isoflurane induced anesthesia). The time series were z-scored over the whole recording session. In order to estimate the dimensionality of each activity space, we estimated the number of principal components that explained more variance than chance.

The time series were split into 5 equal blocks of continuous activity for cross-validation: principal components were estimated using 4 of the blocks (train set) and the remaining block (test set) was projected onto these components to compute the variance they capture : 𝑉𝑎𝑟(𝑋_𝑡𝑒𝑠𝑡_𝑉_𝑡𝑟𝑎𝑖𝑛_𝑉_𝑡𝑟𝑎𝑖𝑛_^T^), with *X_test_* the time series from the test set, and *V_train_* the component matrix of the train set. The procedure was repeated 5 times so that each set was used as the test set once, and the average variance ratio captured in the test sets per component was defined as the cross-validated PC scree plot. A chance level was computed in the same fashion except that the test set was circularly shuffled 20 times in the time dimension independently for each neuron to break temporal relationships between neurons. The 95th percentile of this new variance ratio captured per component distribution was kept as the chance level. The dimensionality of an activity space was defined as the number of components for which the captured variance remains above this chance level, as visualized using scree plots (**Extended Data** Fig. 4a).

#### Spontaneous and sound-evoked activity subspaces comparison

To determine whether spontaneous and sound-evoked activity shared the same dimensions we computed an inclusion index of sound-evoked activity in spontaneous subspace for each recording session. Intuitively, the inclusion index captures how much of the sound-evoked activity variance is captured by the principal components derived from the spontaneous activity. The spontaneous activity was split into 10 blocks of contiguous activity with equal duration. Blocks with an odd-numbered label were used as the reference set for spontaneous activity. We therefore defined dimensions of this reference spontaneous subspace as the ensemble of its first n principal components *V_ref_*, n being the previously computed dimensionality of that subspace. The percentage of variance in another time series *X_source_* that can be explained by dimensions of the reference subspace is 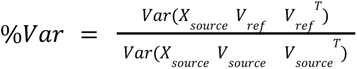 _with *V*_*_source_* the matrix of the first n principal components of *X_source_*. The maximum level of subspace inclusion to be expected *%Var_max_* is obtained when *X_source_* is the other half of spontaneous activity data. The level of inclusion of sound-evoked activity in the spontaneous activity subspace *%Var_projected_*is obtained when *X_source_* is the time series of neuronal activity during sound stimulus presentation (sound-evoked activity). The chance level *%Var_chance_* is calculated with *X_source_* being the sound-evoked activity circularly shuffled in time independently for each neuron. The inclusion index was computed as 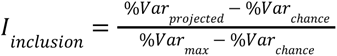.

#### Non-responsive events labelling procedure

To identify sound presentation trials when the neuronal population is not responsive to sounds, we computed cosine similarity of the single-trial sound-evoked response vectors with a template vector computed as the average of the 30% strongest responses to the corresponding sound stimulus. We estimated the variability of that similarity estimator with a bootstrapping procedure : neurons constituting the population were randomly resampled 1000 times with replacement. Trial was labelled as “non-responsive” if 0 belonged in between the 5th and the 95th percentile of the bootstrapped cosine similarity distribution. The analysis was restricted to the 10 sounds driving the strongest responses in the neuronal population for each recording.

#### Quantification and statistical analysis

All quantification and statistical analyses were performed with custom Python scripts. Statistical assessment was based on non-parametric tests reported in the text or in figure legends together with the number of samples used for the test and the nature of the sample (number of neurons, recording sessions or mice). Unless otherwise mentioned, false positive rates below 0.05 were considered significant. In all analyses, all animals who underwent a particular protocol in the study were included.

### Resource availability

**Lead contact**

**Material availability**: Biological material and technologies used in this study are freely available resources.

**Data and code availability**: All datasets and custom codes are freely available on Zenodo. doi:10.5281/zenodo.15530880

## Acknowledgments

We thank Dr. Karim Benchenane for his help in the implementation of delta wave detection. We acknowledge the support of the Fondation pour l’Audition to the Institut de l’Audition.

We acknowledge the support of the following funding sources:

Fondation pour l’Audition, RD-2023-1 (BB), FPA IDA02 (BB) and APA 2016-03 (BB), European Research Council, ERC CoG 770841 DEEPEN (BB)

Fondation pour la Recherche Médicale SPF202005011970 (SB)

Agence Nationale pour la Recherche, AudioDream (BB, SB), France 2030 program, ANR-23-IAHU-0003 (BB)

## Author contributions

Conceptualization: AM, BB, SB Methodology: AM, SB, BB, AF Investigation: AM, SB

Data curation: AM Formal analysis: AM Supervision: BB, SB

Project administration: BB Funding acquisition: BB, SB Software: AM

Visualization: AM Validation: BB, SB

Writing-original draft: BB, SB, AM Writing-review & editing:

## Declaration of interests

The authors declare no competing interests.

## Declaration of generative AI and AI-assisted technologies

The authors did not use generative AI and AI-assisted technologies at any step of the study and in the writing process.

## Supplementary figures

**Extended Data Figure 1:**
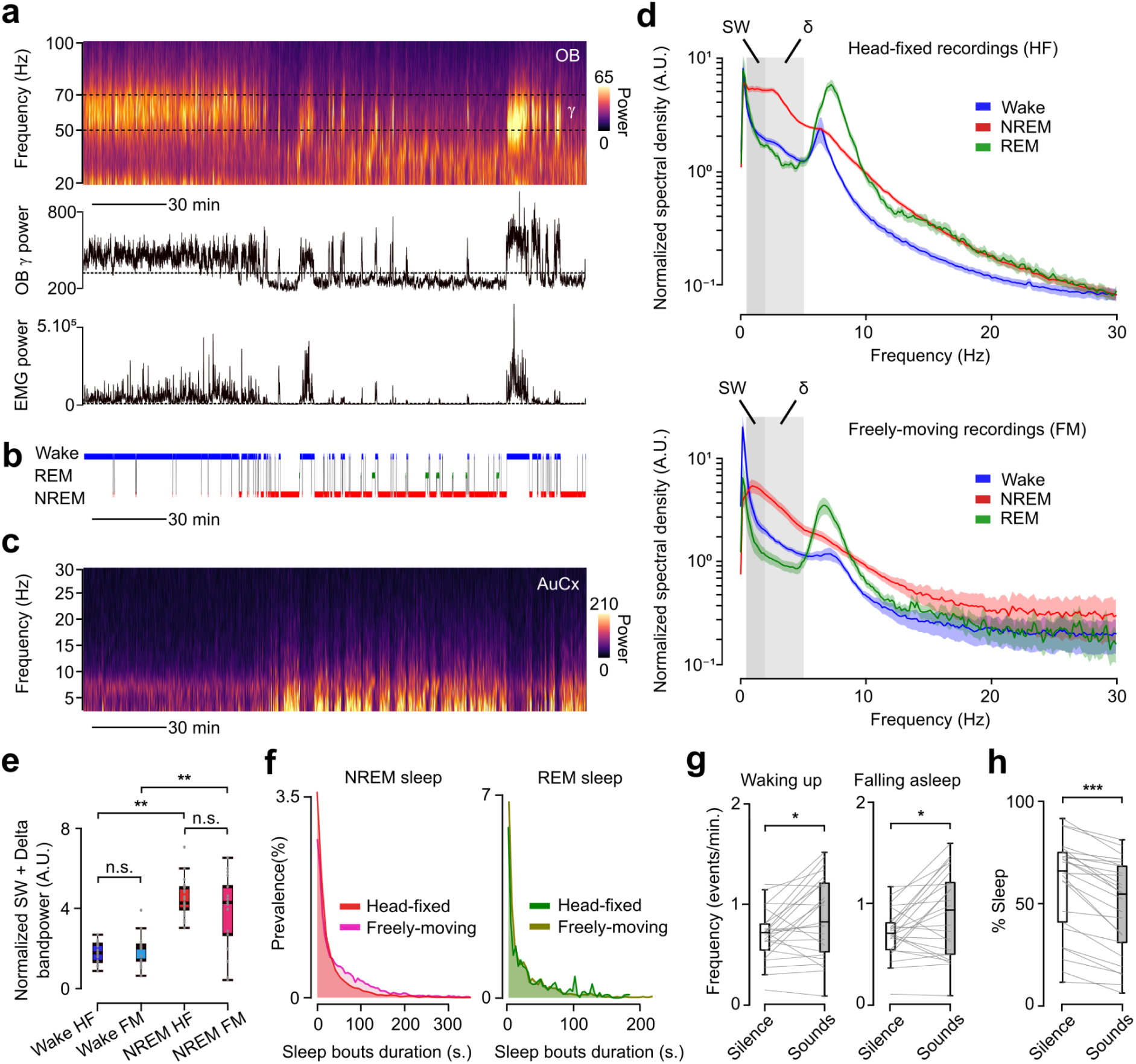
OB gamma and EMG power clearly identify sleep periods in head-fixed mice. **a,** Spectrogram of Olfactory Bulb LFP throughout an entire recording session (top). The evolution of the gamma bandpower (middle) follows EMG power (bottom), and similarly distinguishes sleep, when it is low, from wakefulness, when it is high. **b,** Corresponding hypnogram. **c,** Corresponding auditory cortex spectrogram showing increases in low-frequency power during sleep periods. **d,** Power spectra of auditory cortex LFP during wakefulness, NREM, and REM sleep periods computed with Welch method show similar frequency density when mice were head-fixed (top) or in freely-moving condition (bottom). Power was divided by averaged spectrogram value to allow comparison between recordings. Grey area shows the slow-wave (0.5 - 2 Hz) and Delta (2 - 5 Hz) frequency bands. **e,** Low frequency (0.5 - 5 Hz) power is comparable between head-fixed (HF, n=25 recordings) and freely-moving (FM, n=16 recordings) conditions during wake (Mann-Whitney independent test, : p=8.6×10^−1^) and NREM sleep (Mann-Whitney independent test, p=5.4×10^−1^) and systematically increases in NREM sleep compared to wakefulness (Mann-Whitney independent test, HF : p=1.4×10^−9^ ; FM : p=4.7×10^−3^). **f,** Distribution of sleep bouts duration shows a higher proportion of short NREM sleep periods in head-fixed mice compared to freely-moving but similar REM sleep durations. **g,** Number of wake-up (left) and fall-asleep (right) events slightly increase in “sounds” blocks compared to “blank” blocks (Wilcoxon paired test, wake-up : p=2.0×10^−2^ ; fall-asleep : p=3.4×10^−2^, n=25 recordings). **h,** Percentage of time spent asleep slightly decreases in “sounds” blocks compared to “blank” blocks (Wilcoxon paired test, p=2.5×10^−3^, n=25 recordings).

**Extended Data Figure 2:**
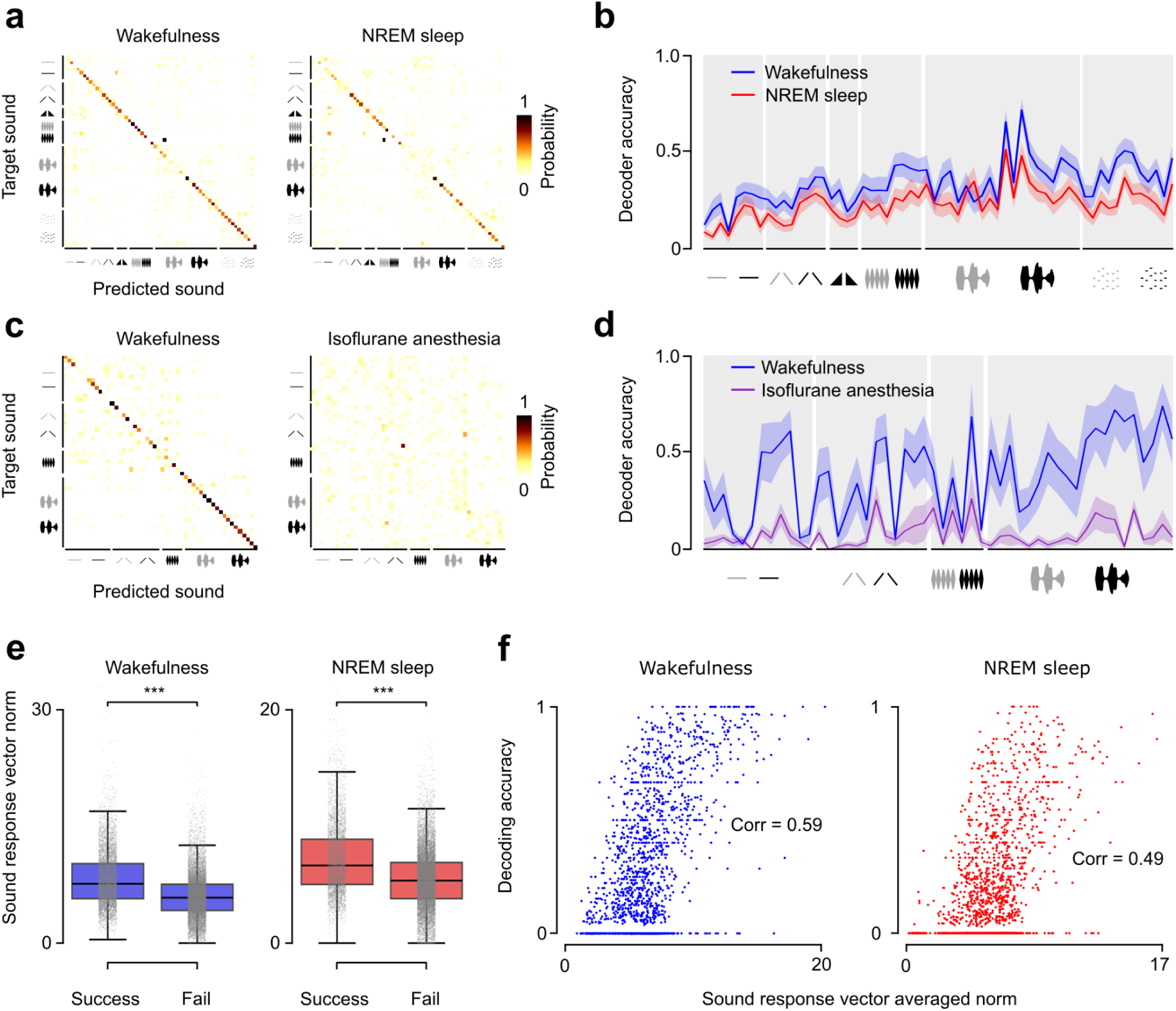
Comparison of sound decoding in different states. **a,** Confusion matrices recapitulating classification outcomes of sound decoders from wakefulness (left) and NREM sleep (right) neuronal activity. **b,** Decoding accuracy for every sound stimulus in wakefulness and NREM sleep. **c,** Confusion matrices recapitulating classification outcomes of sound decoders from wakefulness (left) and isoflurane anesthesia (right) neuronal activity. **d,** Decoding accuracy for every sound stimulus in wakefulness and isoflurane induced anesthesia. **e,** Population response is stronger in trials where the decoder succeeded in classifying sound stimuli both in wakefulness (left, Mann-Whitney rank test, p = 0.0) and NREM sleep (right, Mann-Whitney rank test, p=7.3×10^−241^). **f,** Decoding accuracy of a sound positively correlates with the averaged response intensity of that sound in both wakefulness (p = 2.6×10^−141^) and NREM sleep. (p = 8.6×10^−89^).

**Extended Data Figure 3:**
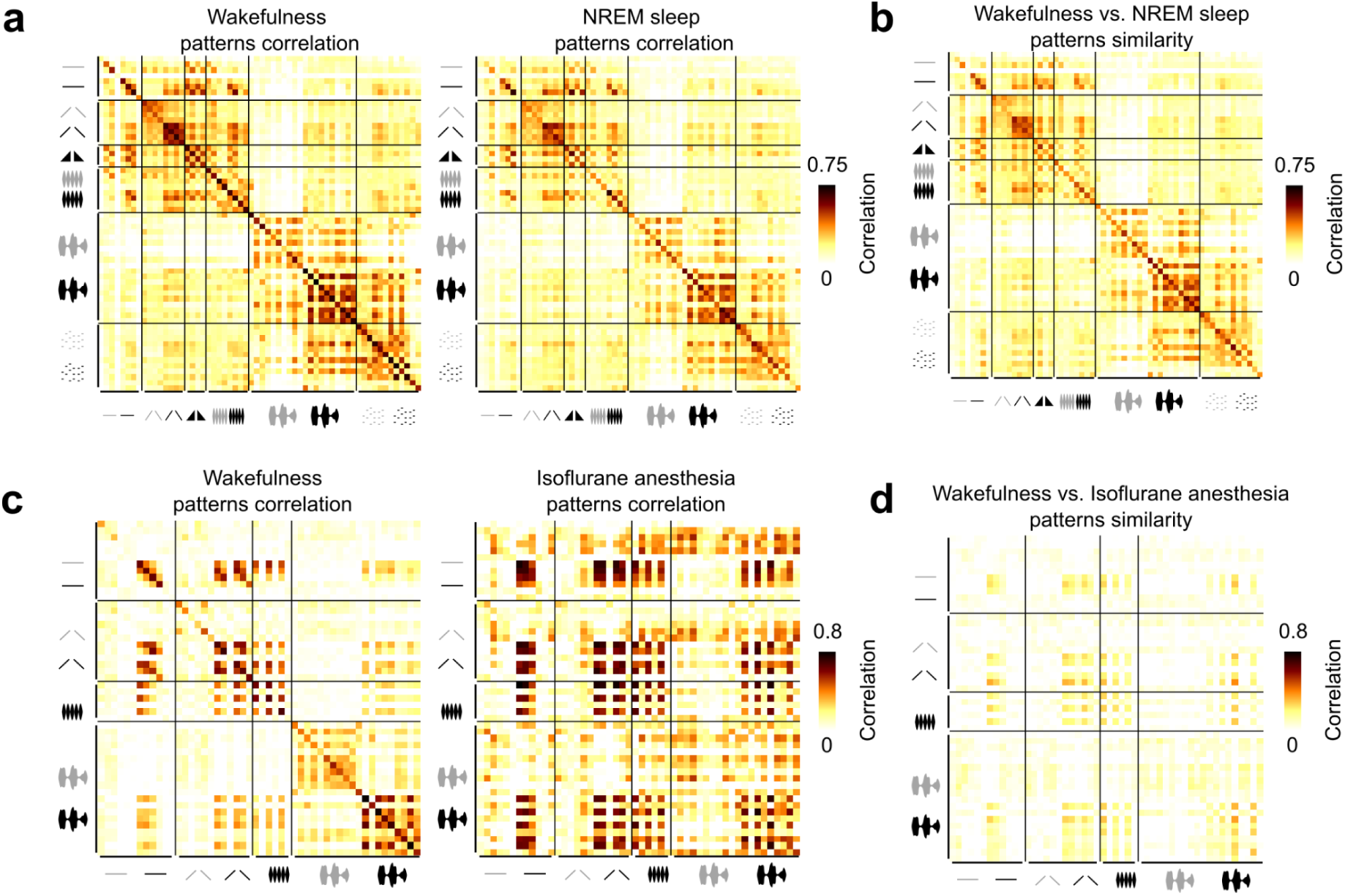
Within-state and cross-state representation similarity analysis: NREM sleep vs wake and anesthesia vs wake. **a,** Non noise corrected similarity matrices between population sound-evoked responses averaged over two distinct halves of the sound-presentations in wakefulness (left), and in NREM sleep (right). **b,** Non noise corrected similarity matrices between sound-evoked responses in wakefulness and NREM sleep. **c,** Same as **a** but for wakefulness and isoflurane anesthesia. **d,** Same as **b** for wakefulness and isoflurane anesthesia.

**Extended Data Figure 4:**
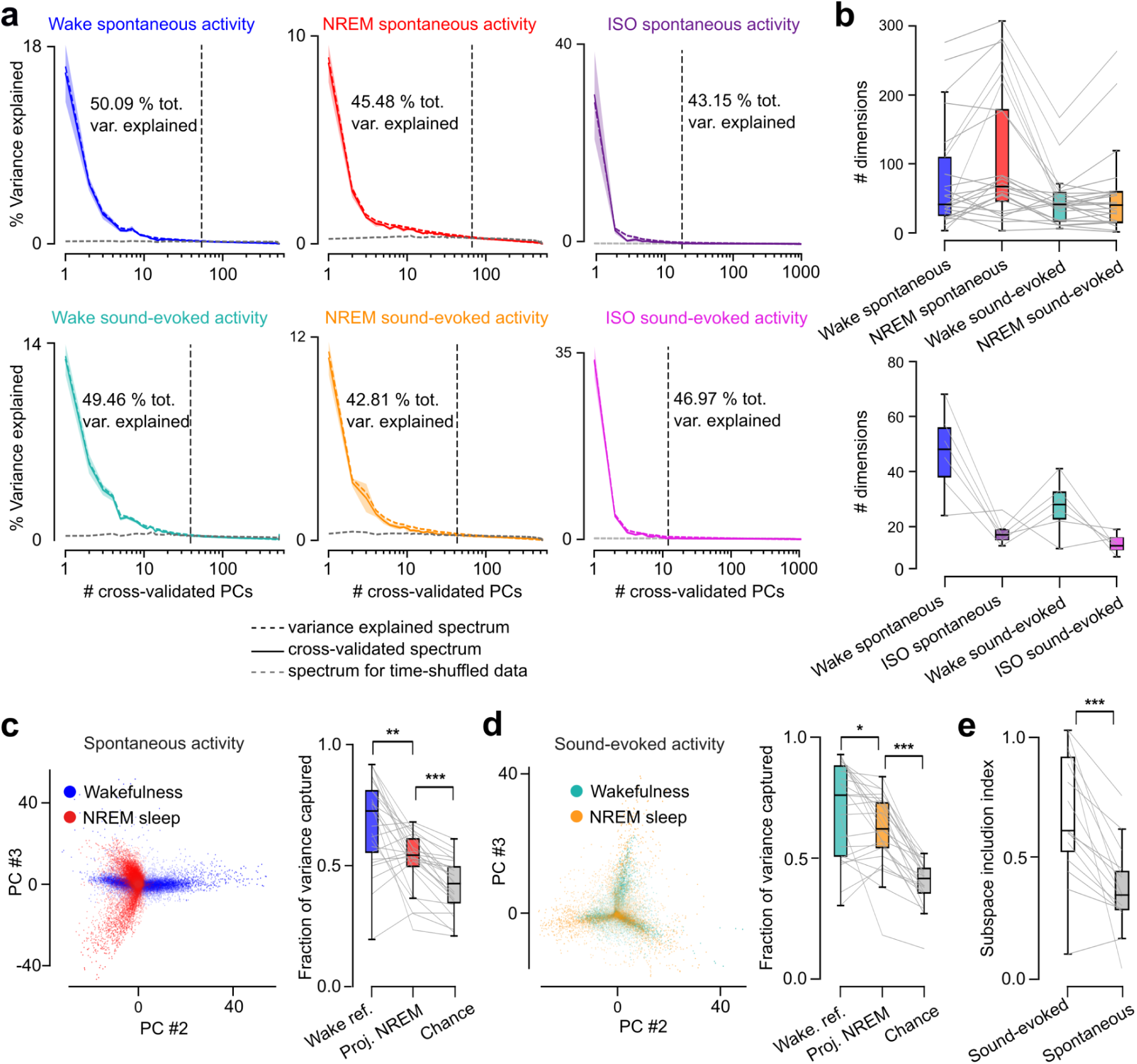
Geometry of spontaneous and evoked activity across sleep and wake. **a,** Scree plots from PCA (dashed) and cross-validated PCA (solid line, see methods) performed on wake, NREM, and isoflurane anesthesia, spontaneous and sound-evoked neuronal activity. Note that the two curves strongly overlap. The number of dimensions used to describe each subspace was taken as the number of principal components (coloured) that capture more variance than expected by chance (grey), and is here denoted as a vertical grey dashed line. About 50% of total variance is usually captured in that subspace. **b,** Number of components used to describe each condition subspace. **c,** Left : 2D example projection of wakefulness and NREM sleep spontaneous neuronal activity in principal component (PC) space. Right : Fraction of variance of spontaneous wakefulness and NREM sleep neuronal activity explained by dimensions of the spontaneous wakefulness subspace (top, Wilcoxon paired test, “Wake reference” to “Projected NREM” : p=2.5×10^−3^; “Projected NREM” to “chance level” : p=6.0×10^−8^, n=25). **d,** Same as **c** for sound-evoked activity (Wilcoxon paired test, “Wake reference” to “Projected NREM” : p=2.7×10^−2^; “Projected NREM” to “chance level” : p=6.0×10^−8^, n=25). **e,** The inclusion index of NREM sleep sound-evoked activity in wake sound-evoked subspace is higher than the inclusion index of spontaneous NREM sleep spontaneous activity in wake spontaneous subspace (Wilcoxon paired test, p=1.2×10^−4^, n=15). Sessions where the maximum overlap expected value did not exceed chance level by 0.1 were removed.

**Extended Data Figure 5 :**
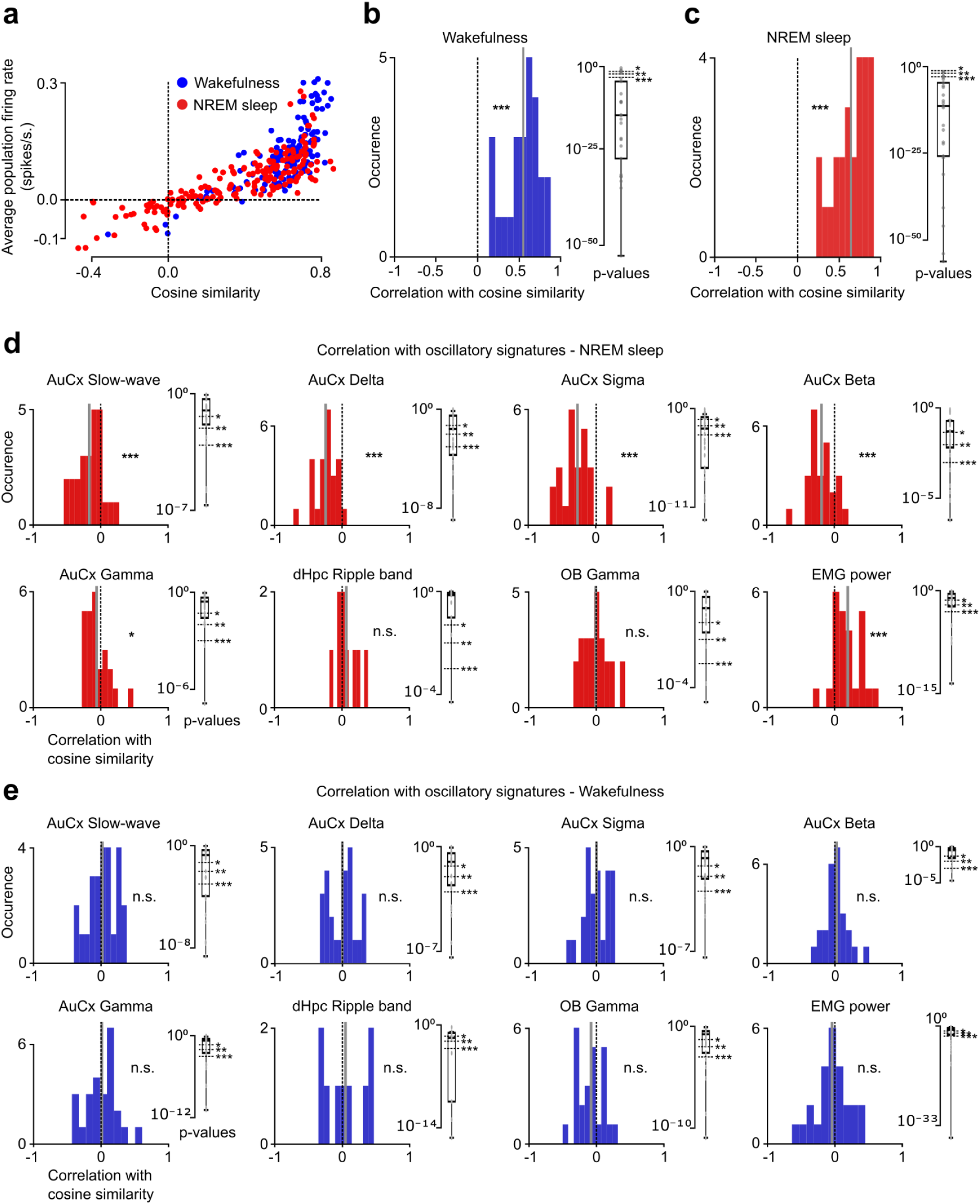
Sound encoding reduction is explained by population response gain, and increases with Delta and Sigma oscillatory power in NREM sleep but not in wake. **a,** Example of positive relationship between amplitude of population responses to sounds and their similarity to the expected response vector in wakefulness and NREM sleep from a single recording. **b,** Left: distribution of Pearson correlation coefficients between the amplitude of population responses to sounds and their similarity to the expected response vector in wakefulness for every recording (Wilcoxon signed rank test, p = 6.0×10^−8^, n = 25). The solid grey vertical line indicates the mean of the distribution. Right: correlation p-values. Dashed lines indicate common values to assess statistical significance (* = 0.05, ** = 0.01, *** = 0.001). **c,** Same as **b** in NREM sleep (left, Wilcoxon signed-rank test, p = 6.0,10^−8^, n = 25). **d,** Distribution of correlations coefficients between response similarity level and bandpower in the contralateral AuCx LFP in NREM sleep (Wilcoxon signed rank, Slow-wave : p=2.9×10^−4^ ; Delta : p=4.2×10^−7^ ; Sigma : p=1.8×10^−5^ ; Beta ; p=3.2×10^−5^; Gamma : p=3.2×10^−2^, n=25 recordings), the ripple bandpower in the dorsal Hippocampus (Wilcoxon, paired test, p=5.7×10^−1^, n=8), and the OB gamma band and filtered EMG power (Wilcoxon paired test, OB Gamma : p=8.1×10^−1^ ; EMG : p=4.5×10^−5^, n=25 recordings). Pearson correlation p-values for single recordings are shown on the right. **e,** Same as **d** in wakefulness (Wilcoxon signed rank test, AuCx SW : p=6.0×10^−1^; AuCx Delta : p=9.8×10^−1^; AuCx Sigma : p=9.4×10^−1^; AuCx Beta : p=4.7×10^−1^; AuCx Gamma : p=7.5×10^−1^ n=25; dHpc ripple band : p=7.3×10^−1^, n=8; OB Gamma : p=8.5×10^−2^; EMG power : p=4.9×10^−1^, n = 25)

**Extended Data Figure 6:**
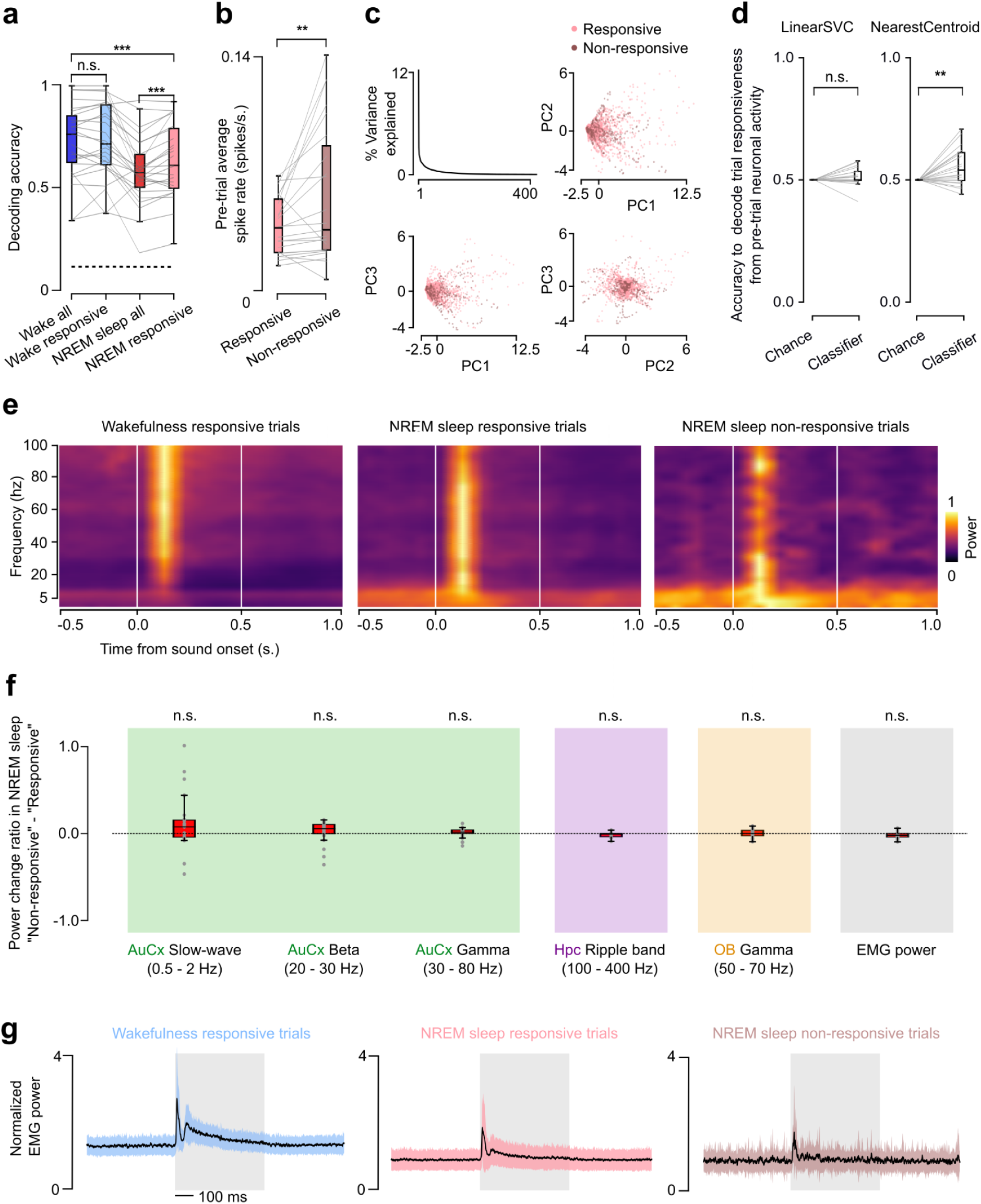
Population gating events are correlated to brain activity. **a,** Sound decoding accuracy during wakefulness is unchanged when “non-responsive” events are removed (Wilcoxon paired test, p=6.1×10^−1^, n=25 recordings), but slightly increased in NREM sleep (Wilcoxon paired test, p=4.9×10^−4^, n=25 recordings). Removing “non-responsive” events from NREM sleep data was not sufficient to reach the accuracy level of wakefulness data (Wilcoxon paired test, p=1.2×10^−4^, n=25 recordings). **b,** Averaged neuronal activity in the 500 ms preceding a “non-responsive” shows increased activity compared to “responsive” event in NREM sleep (Wilcoxon paired test, p=1.3×10^−3^, n=25 recordings). **c,** PCA on neuronal activity in the 500 ms preceding sound presentation during sleep. Top left : scree plot. Top right and bottom : 2D example projections show no subspace distinction between activity preceding a “responsive” and a “non-responsive” event. **d,** LinearSV decoder trained to classify “responsive” and “non-responsive” events from the neuronal activity preceding sound presentation is unable to perform over chance level (Wilcoxon paired test, p=1.2×10^−1^, n=25 recordings), but a Nearest Centroid classifier performs better than chance (Wilcoxon paired test, p=7.2×10^−3^, n=25 recordings). Chance level was calculated by randomly shuffling the classification labels. **e,** Averaged spectrograms of contralateral auditory cortex LFP around sound presentation in “responsive” events in wakefulness, “responsive” and “non-responsive” events in NREM sleep. Each frequency band was divided by its maximum value from the three spectrograms to witness changes in time and allow comparison between conditions. **f,** Relative change from “responsive” to “non-responsive” events in oscillation power in a 1.5 s long window centered around sound presentation, positive values indicate increases in “non-responsive” compared to “responsive” events. Slow-wave, Beta, and Gamma oscillation power in the contralateral AuCx are unchanged in “non-responsive” compared to “responsive” events (Wilcoxon signed rank test, Slow-wave : p=5.2×10^−2^ ; Beta : p=9.8×10^−2^ ; Gamma : p=1.3×10^−1^, n=25 recordings), as well as the ripple bandpower in the dorsal Hippocampus (Wilcoxon signed rank test, p=4.7×10^−1^, n=8), the Olfactory Bulb gamma power (Wilcoxon signed rank, p=6.2×10^−1^, n=25), and the filtered EMG power (Wilcoxon signed rank test, p=8.0×10^−2^, n=25 recordings). **g,** Filtered EMG power around sound presentation shows a peak at sound onset in both wakefulness and NREM sleep regardless of the auditory cortex neurons population being responsive or not. Standard deviation is displayed as a colored area.

**Extended Data Figure 7:**
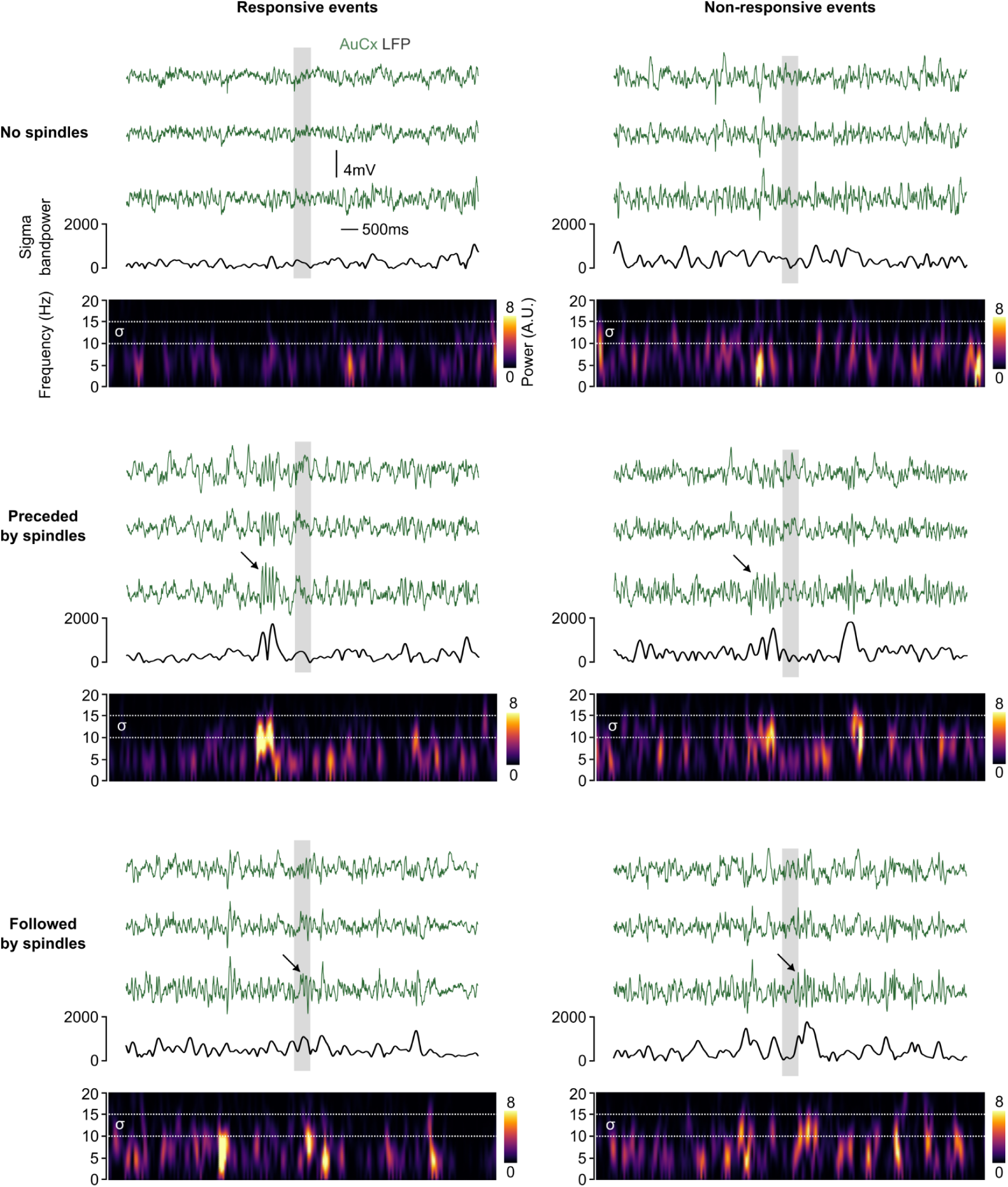
Responsiveness of neuronal population to sounds does not depend on the occurrence of sleep spindles. LFP recording of the contralateral auditory cortex shows that “responsive” (left) and “non-responsive” (right) events can both be surrounded or not by sleep spindles. Filtered AuCx LFP traces (0.5 - 70 Hz) from 3 different channels are shown in green. The sigma (10-15 Hz) power of the bottom trace is shown in black with its spectrogram. Arrows indicate a visually detected sleep spindle event. Sound presentation is shown as a grey area.

**Extended Data Figure 8:**
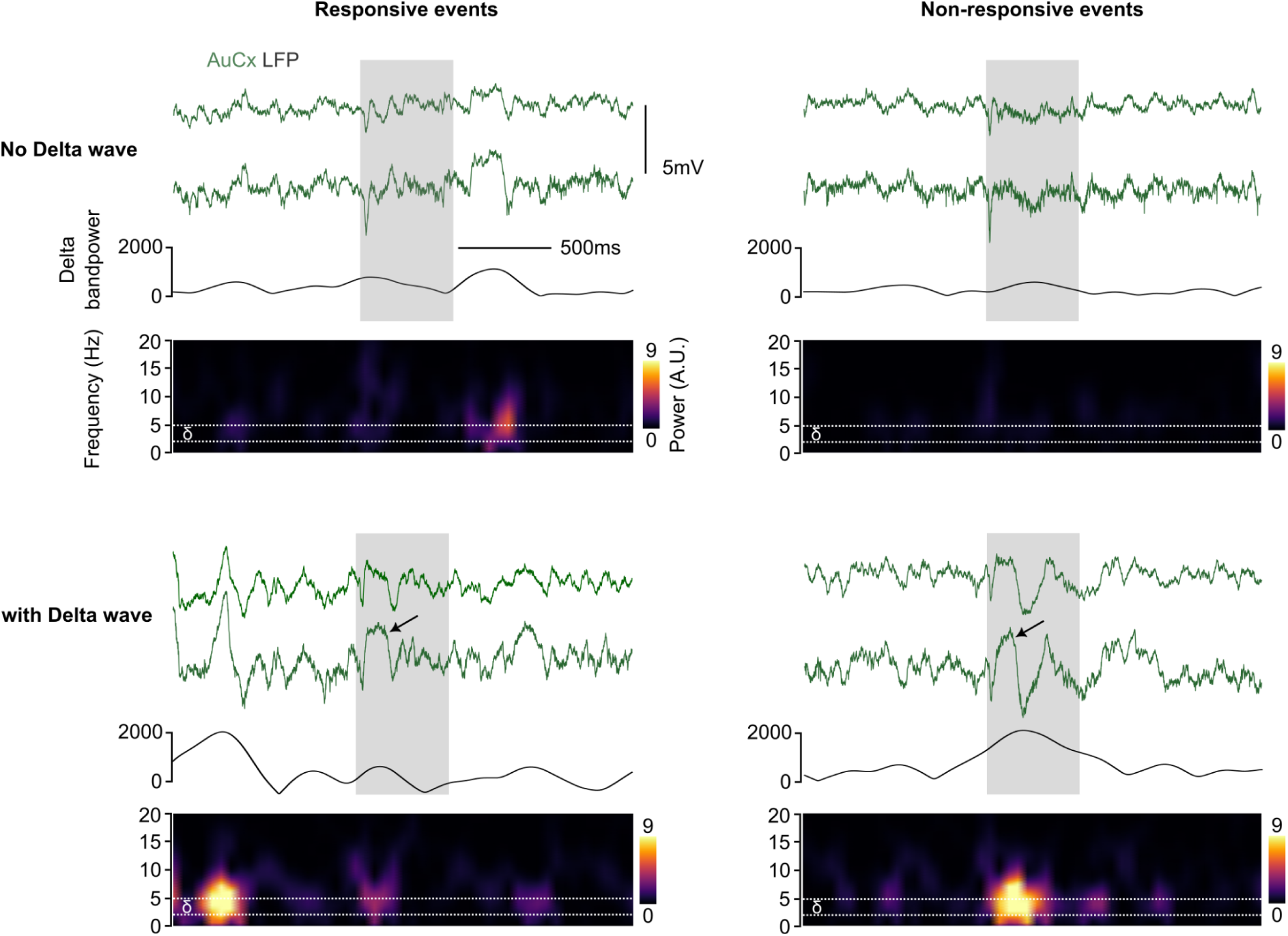
Responsiveness of neuronal population to sounds does not depend on the occurrence of delta waves. LFP recording of the contralateral auditory cortex shows that “responsive” (left) and “non-responsive” (right) events can both co-occur or not with a delta wave. AuCx traces from 2 different channels are shown in green. The delta (2-5 Hz) power of the bottom trace is shown in black with its spectrogram. Arrows indicate a visually detected delta wave event during the sound presentation. Sound presentation is shown as a grey area.

## Notes

doi:10.5281/zenodo.15530880

### Competing Interest Statement

The authors have declared no competing interest.

